# LRG1 promotes atherosclerosis by activating macrophages

**DOI:** 10.1101/2024.01.23.576507

**Authors:** Juan Wang, Sitao Zhang, Jing Wang, Jiuchang Zhong, Hongbin Liu, Weiming Li, Mulei Chen, Li Xu, Wenbin Zhang, Ze Zhang, Zhizhong Wei, Jia Guo, Xinyu Wang, Jianhua Sui, Xingpeng Liu, Xiaodong Wang

## Abstract

**Background:** Atherosclerosis is a chronic inflammatory disease of the arterial wall characterized by the accumulation of cholesterol-rich lipoproteins in macrophages. Leucine-rich alpha-2 glycoprotein 1 (LRG1) is a circulating protein associated with inflammation, however, its role in atherosclerosis remains unclear. This study identified its role in macrophage pro-inflammatory differentiation and revealed the relationship between LRG1 and atherosclerosis.

**Method:** We evaluated the impact of LRG1 on atherosclerosis progression by analyzing atherosclerotic tissue and serum samples from patients with coronary artery disease (CAD) and healthy individuals and analyzed its role in such a process using two types of mice models: *Apoe* knock-out mice (*Apoe^-/-^*) and *Apoe* and *Lrg1* double knock-out mice (*Apoe^-/-^/Lrg1^-/-^*). These mice were fed with a high-fat diet for 16 to 32 weeks to simulate conditions exacerbating atherosclerosis. To examine the effects of inhibiting LRG1 on atherogenesis, we administered intraperitoneal injections of LRG1 neutralizing antibody (50μg/kg) weekly to *Apoe^-/-^* mice for 8 weeks. We conducted *in vitro* assays using bone marrow-derived macrophages isolated from wild-type mice and analyzed transcriptional signatures using RNA sequencing. Additionally, we utilized small molecular inhibitors to validate the signaling pathway through which LRG1 promotes macrophage-driven inflammation.

**Results:** LRG1 levels were found to be elevated in patients with atherosclerosis and correlated with higher levels of a plasma pro-inflammatory biomarker high-sensitive C-reactive protein (hsCRP), and several macrophage-related pro-inflammatory markers including CD68, VE-Cadherin and VCAM-1. In a high fat diet induced *Apoe^-/-^* mouse atherosclerosis model, the deletion of *LRG1* gene significantly delayed atherogenesis progression and reduced levels of macrophage-related pro-inflammatory cytokines. Addition of purified LRG1 to cultured macrophages stimulated those macrophages to pro-inflammatory M1-like polarization regulated by the activation of ERK and JNK pathways. An anti-LRG1 neutralizing antibody effectively blocked LRG1-induced macrophage M1-like polarization *in vitro* and conferred therapeutic benefits to animals with ApoE deficiency-induced atherosclerosis.

**Conclusion:** LRG1 plays an important pro-inflammatory role in atherosclerosis by influencing macrophage polarization towards a pro-inflammatory state. The inhibition of LRG1 with neutralizing antibodies may offer a potential therapeutic strategy for patients with atherosclerosis by mitigating the pro-inflammatory response and delaying disease progression, offering a novel therapy in atherosclerosis management.

**Translational Perspective:** Atherosclerosis, a persistent inflammatory condition affecting the arterial wall, serves as the underlying pathophysiological basis for acute ischemic cardiovascular events. The involvement of macrophages is crucial in the advancement of atherosclerosis. In this investigation, heightened levels of plasma LRG1 were observed in individuals with coronary artery disease. Moreover, this study presents initial evidence highlighting LRG1 as a pivotal activator of macrophages, instigating a pro-inflammatory M1 polarization during atherogenesis through the activation of ERK1/2 and JNK pathways. The use of an anti-LRG1 neutralizing antibody demonstrated a delay in atherosclerosis progression in an animal model, suggesting a potential therapeutic target for atherosclerosis treatment. Suppression of LRG1 production could impede atherosclerosis advancement and enhance plaque stability. Utilizing neutralizing antibodies against LRG1 emerges as a promising therapeutic approach for treating atherosclerosis.

## Introduction

Atherosclerosis a chronic inflammatory disorder of the arterial wall, wherein modified lipoproteins enriched with cholesterol become entrapped within the subendothelial region of the vessel wall.^1^ This event triggers the recruitment of inflammatory monocytes from the circulatory system, leading to immune responses that significantly contribute to the onset, progression, and clinical complications of atherosclerosis.^2,3,4^ Lesional macrophages emerge through the recruitment of circulating monocytes, as well as local proliferation.^2,5^ Particularly, the inflamed macrophage assumes a pivotal role in driving the advancement and evolution of atherosclerosis, with its pro-inflammatory activities potentially fostering instability and rupture of arterial plaques.^6^ The accumulation of lipid-laden macrophages exhibiting a “foamy” morphology constitutes a hallmark of atherosclerotic lesions.^7^ A central inquiry in the field pertains to the mechanisms governing the differentiation and activation of macrophages during the development of coronary artery disease (CAD).^8^

Leucine-rich alpha-2 glycoprotein 1 (LRG1) is a circulating protein belonging to the highly conserved leucine-rich repeat (LRR) protein family.^8^ It is primarily synthesized and secreted by the liver and immune cells, serving as an acute-phase protein.^9^ LRG1 exhibits elevated expression in numerous inflammatory conditions, including rheumatoid arthritis, pulmonary disorders, diabetic kidney disease, heart failure, and microbial infection.^10–17^ The elevated plasma levels of LRG1 seems to correlate with peripheral artery disease, along with various cardiovascular risk factors such as arterial stiffness, endothelial dysfunction and obesity.^18^ However, the precise role of LRG1 in atherogenesis remains unclear, and its causal impact on atherosclerosis has yet to be elucidated. The current study suggested the causative role of LRG1 in atherosclerotic inflammation. The research first revealed the pronounced LRG1 expression within atherosclerotic lesions in both human subjects and mice. Notably, the absence of LRG1 attenuates atherosclerosis progression, promoting plaque stability characterized by reduced necrosis and thicker fibrous caps, along with a reduction in circulating pro-inflammatory monocytes. *In vitro* experiments demonstrate that LRG1 fosters the polarization of macrophages toward a M1-like phenotype, resulting in the upregulation of pro-inflammatory gene expression in these cells. The neutralization of LRG1 by an anti-LRG1 antibody inhibited the M1-like macrophage activation and led to a significant reduction in aortic lesion area. Remarkably, RNA-sequencing analysis of LRG1-treated macrophages reveals the amplification of the inflammatory response, marked by heightened expression of pro-inflammatory genes. These findings firmly establish the role of LRG1 as a secreted pro-atherosclerotic protein, uncovering the mechanisms governing its impact on macrophage inflammation, monocyte migration, and atherosclerosis. Collectively, this research indicates that LRG1 initiates inflammation in atherosclerosis, offering a novel target for preventive therapeutic intervention in the treatment of this condition.

### Methods

Detailed methods are provided in the Supplemental Material. The authors declare that all supporting data are available within the article (and its Supplemental Material). The study protocol was approved by the Ethics Committee of Beijing Chao-Yang Hospital and Chinese PLA General Hospital and written informed consent was obtained from all patients. Human samples used in this study are fully described in Supplemental Methods. Human femoral arteries were obtained from surgical vascular grafting operations.

### Statistical analysis

We present continuous variables as mean and standard deviation (SD) or standard error of the mean (SEM), as specified, and summarize categorical data as frequencies or percentages. We evaluated the existence of a normal distribution with the Kolmolgorov-Smirnov test. An unpaired 2-sided Student t test was used when data were in normal distribution. A nonparametric test (Mann-Whitney) was used when data did not pass the normality test or sample size was small. We analyzed differences among patient groups by one-way analysis of the Kruskal-Wallis analysis, followed by post hoc analysis with the Bonferroni’s correction. We determined correlation between variables by the non-parametric tests (Spearman rank order correlations, r_s_) and p-value< 0.05 was considered statistically significant. We constructed two models for multivariable logistic regression to assess the independent determinants of CAD. In Model 1, we included traditional risk factors. In Model 2, we adjusted the analysis for the LRG1. We performed receiver operator characteristic (ROC) analysis of risk factors and biomarkers measurements with and without LRG1. For cell or animal experiments, data represent the average of six experiments. Data analysis was performed using GraphPad Prism Software Version 9 (GraphPad). For Bulk RNA-seq, the Differential Gene Expression-Gene Set Analysis algorithm was implemented. All analyses used two-sided tests with an overall significance level (α) of 0.05, and were performed with the SPSS 15.0 for Windows (SPSS, Inc., Chicago, IL, USA) and SAS Version 9.1 (SAS Institute, Cary, NC, USA) software.

## Results

### The elevated levels of LRG1 in the plasma of patients with CAD are associated with biomarkers of CAD

To explore the potential correlation between LRG1 and the progression of CAD, we conducted an analysis of 30 traditional cardiovascular risk factors and biomarkers from 318 CAD patients (patients’ source) and 122 non-CAD individuals (Table 1). Among the 30 variables we analyzed, we observed that plasma LRG1 levels were significantly higher in CAD patients compared to non-CAD individuals (Figure 1A). Furthermore, we identified a statistically significant correlation between LRG1 and multiple inflammation biomarkers (Table 2), including hsCRP (r_s_ = 0.279, P < 0.001), growth differentiation factor-15 (GDF-15) (r_s_ = 0.148, P < 0.002), and fibrinogen (r_s_ = 0.246, P < 0.001). These findings suggest that the serum level of LRG1 is associated with CAD and may play a role in the inflammatory processes during atherosclerosis progression.

**Figure 1.**
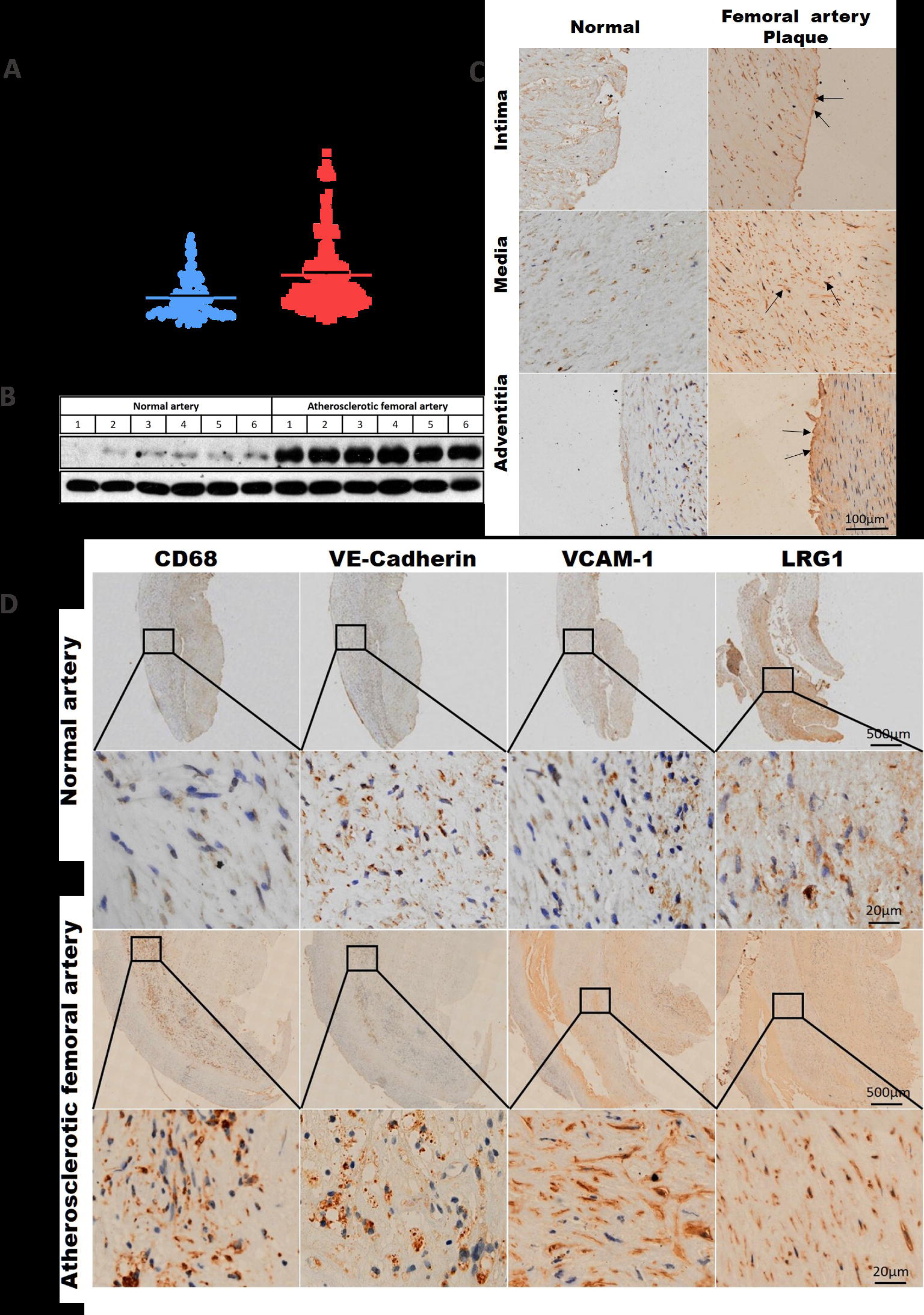
LRG1 elevation correlates with heightened macrophage-derived proinflammatory cytokines in femoral plaques of atherosclerotic femoral stenosis patients. **(A)** ELISA analyses the plasma LRG1 levels of age-matched CAD and non-CAD patients. **(B)** Immunoblotting detects the LRG1 levels of samples from atherosclerotic femoral arteries of 10 patients with severe femoral stenosis. Non-femoral stenosis arteries are obtained from surplus-otherwise discarded segments of non-atherosclerotic femoral arteries during surgical vascular grafting operations and frozen immediately in liquid nitrogen. GAPDH is shown as the sample loading control. **(C)** The immunohistochemical staining of LRG1 in the intima, media, and adventitial regions of atherosclerotic or normal arteries slides from CAD patients. **(D)** The immunohistochemical staining of LRG1 and chronic atherosclerosis markers (CD68, VE-Cadherin, and VCAM-1) of femoral stenosis or normal arteries slides from femoral stenosis patients. [Mean ± SEM. n.s., not significant, *P < 0.05, **p<0.01, ***p<0.001; One-way ANOVA followed by Tukey’s multiple comparison test]

**Table 1.**
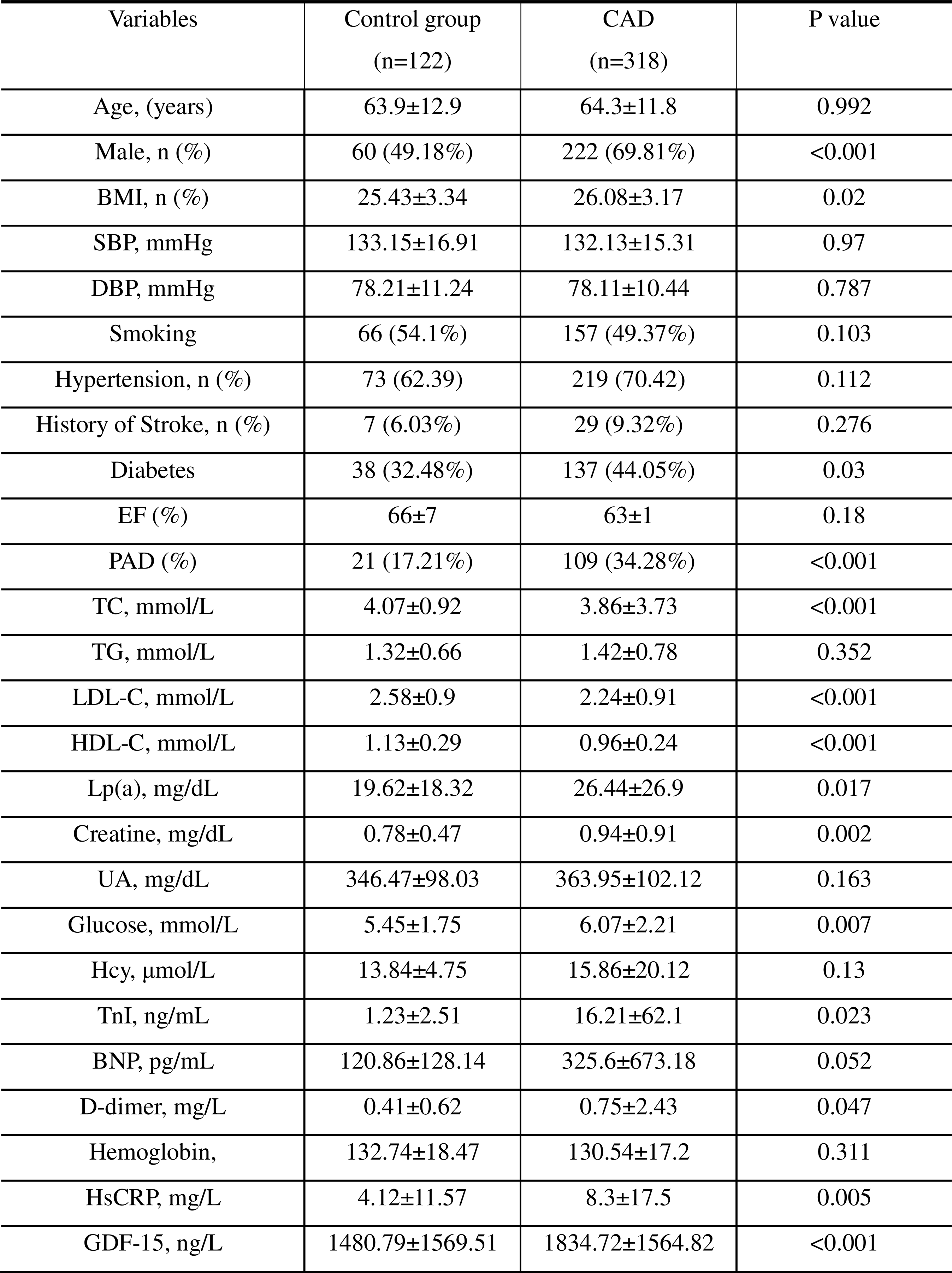
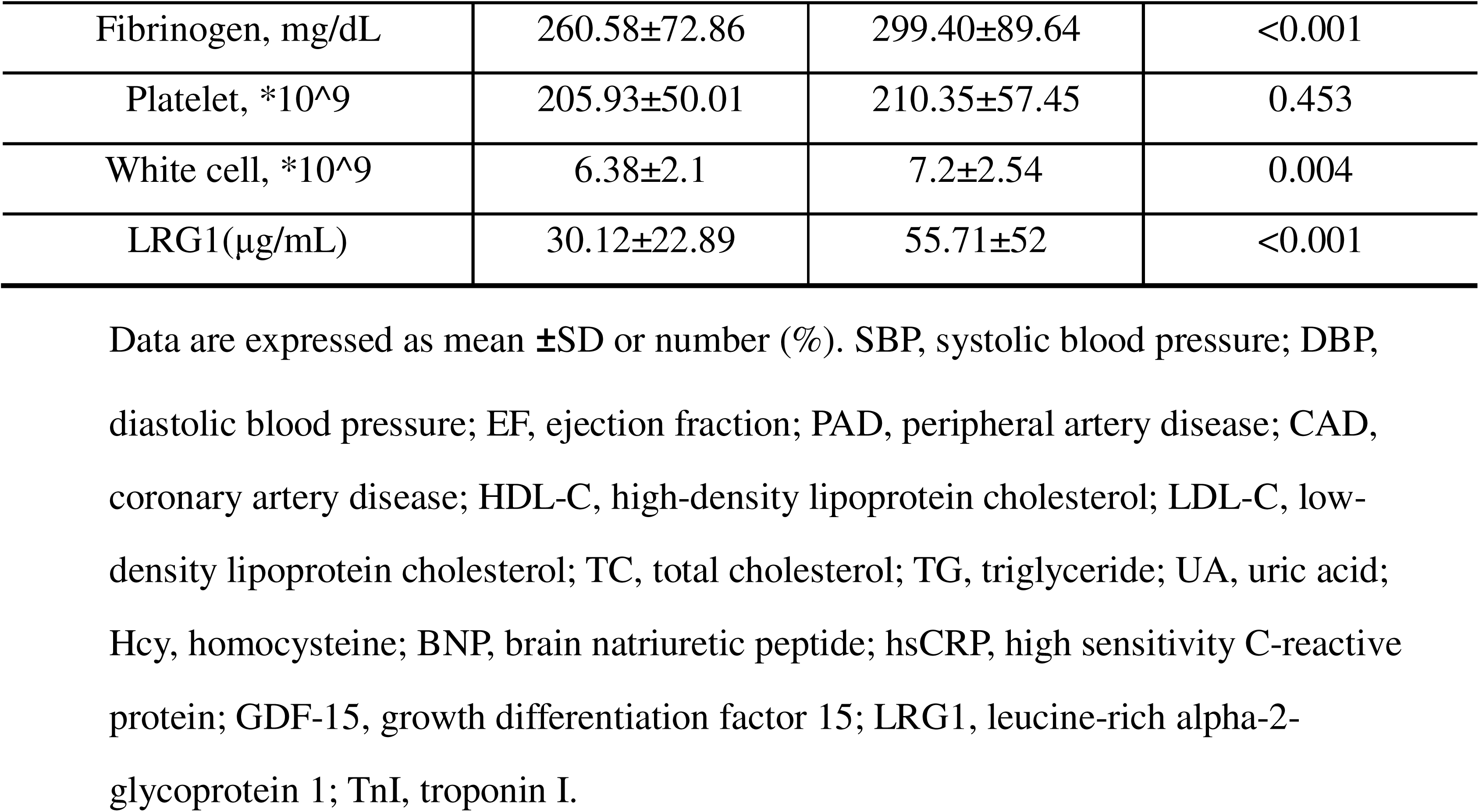
Baseline characteristics and cardiovascular risk factors of both CAD and non-CAD patients.

**Table 2.**
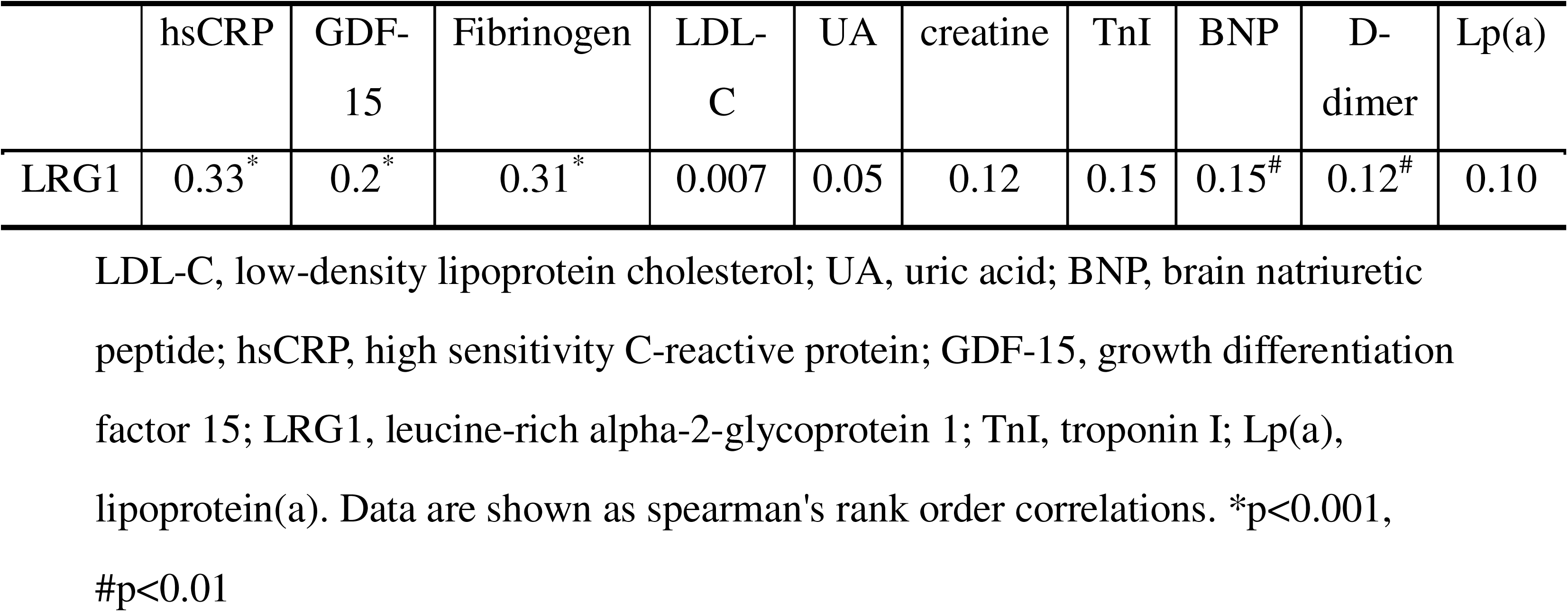
Correlation between LRG1 with plasma inflammatory biomarkers.

### Verification of LRG1 as an independent biomarker for CAD

LRG1 has long been recognized as a biomarker for diabetes and in high-cardiovascular risk scenarios.^10^ To further authenticate the potential of LRG1 as an independent biomarker for CAD, we conducted comprehensive multivariable analyses encompassing 13 traditional cardiovascular risk factors (as delineated in Table 1). In the initial Model 1, various variables such as age, gender, body mass index (BMI), smoking habits, hypertension, diabetes, history of stroke, Low-Density Lipoprotein Cholesterol (LDL-C), High-Density Lipoprotein Cholesterol (HDL-C), Total Cholesterol (TC), Lipoprotein(a) [Lp(a)], creatinine, and Troponin I (TnI) were found to be associated with CAD (Table 3). Upon the inclusion of LRG1 in the analysis in Model 2, LRG1 levels continued to emerge as an independently associated factor with CAD, retaining its significance alongside the conventional elements and biomarkers included in Model 1 (as depicted in Table 3). This substantiates the proposition that LRG1 serves as a distinct and self-reliant biomarker for individuals afflicted by CAD in the realm of clinical practice.

**Table 3.** Multivariable regression analysis of independent risk factors for coronary artery disease.

| variable | OR (95% CI) | P-value |
| --- | --- | --- |
| Model 1 |  |  |
| Age | 1.0 (0.98, 1.02) | 0.86 |
| Sex | 1.45 (0.83, 2.52) | 0.19 |
| BMI | 1.02 (0.94, 1.1) | 0.64 |
| Hypertension | 1.48 (0.83, 2.65) | 0.18 |
| Smoking | 1.0 (0.98, 1.02) | 0.86 |
| Stroke | 1.1 (0.4, 3.04) | 0.86 |
| Diabetes | 1.29 (0.72, 2.3) | 0.4 |
| LDL-C | 0.53 (0.15, 1.89) | 0.33 |
| HDL-C | 0.12 (0.02, 0.89) | 0.038 |
| TC | 1.5 (0.34, 6.5) | 0.59 |
| Lp (a) | 1.02 (1-1.03) | 0.014 |
| Creatine | 1.03 (0.7, 1.51) | 0.87 |
| TnI | 1.03 (0.98, 1.07) | 0.27 |
| Model 2 |  |  |
| Age | 0.99 (0.97, 1.02) | 0.613 |
| Sex | 1.74 (0.96, 3.16) | 0.068 |
| BMI | 1.04 (0.95, 1.13) | 0.429 |
| Hypertension | 1.51 (0.82, 2.78) | 0.19 |
| Smoking | 0.91 (0.52-1.57) | 0.73 |
| Stroke | 1.09 (0.36-3.29) | 0.875 |
| Diabetes | 1.63 (0.88-3.03) | 0.12 |
| LDL-C | 0.45 (0.12-1.72) | 0.245 |
| HDL-C | 0.15 (0.02-1.22) | 0.076 |
| TC | 1.96 (0.29-13.37) | 0.467 |
| Lp (a) | 1.02 (1.0, 1.04) | 0.89 |
| Creatine | 0. (0.6, 1.34) | 0.604 |
| TnI | 1.03 (0.98, 1.08) | 0.256 |
| LRG1 | 1.03 (1.02, 1.04) | <0.001 |

Data are represented as odds ratio (95% confidence interval). OR, Odds Ratio, CI, confidence interval; HDL-C, high-density lipoprotein cholesterol; LDL-C, low- density lipoprotein cholesterol; TC, total cholesterol; LRG1, leucine-rich alpha-2- glycoprotein 1; TnI, troponin I; Lp(a), lipoprotein(a). Two models were used in the analysis. Model 1 was adjusted for 13 conventional cardiovascular risk factors, while Model 2 included these 13 factors from Model 1 and further incorporated LRG1 as an additional variable.

### LRG1 elevation correlates with increased macrophage-derived proinflammatory cytokines in femoral plaques of atherosclerotic femoral stenosis patients

To further explore the potential correlation between LRG1 levels and the progression of atherosclerosis, we collected atherosclerotic femoral arteries from 10 patients with severe femoral stenosis during the surgical vascular grafting operations. Non-femoral stenosis arteries were obtained from surplus segments of non-atherosclerotic femoral and used as controls for comparison with the atherosclerotic femoral arteries. Through immunoblotting analysis, we observed a significant increase in LRG1 protein levels in all 6 femoral plaque samples compared to the 6 normal arteries (Figure 1B). Additionally, immunohistochemical analysis of atherosclerotic femoral arteries slides from CAD patients also revealed a significant increase in LRG1 signal in the intima, media, and adventitial regions of the femoral plaques when compared to the corresponding regions of normal arteries (Figure 1C, the relative quantifications shown in Figure S1A-C). These findings suggest that LRG1 is associated with the formation of femoral plaque and the progression of atherosclerosis. In Table 2, we made a noteworthy observation of a positive correlation between elevated plasma LRG1 levels and multiple inflammatory biomarkers associated with CAD. To further validate whether increased LRG1 levels are also linked to chronic inflammatory markers related to atherosclerotic lesions, we conducted immunohistochemical staining on femoral plaque samples and healthy arteries (used as controls) obtained from the same femoral stenosis patients. Our findings revealed significantly higher expression levels of three inflammatory-related biomarkers, namely CD68, VE-Cadherin, and VCAM-1, in the femoral plaque samples compared to the healthy arteries from the same donors (as shown in Figure 1D, the relative quantifications shown in Figure S1D-G). Moreover, during the detection of LRG1 levels using continuous slides, regions with accumulated signals of CD68, VE-Cadherin, and VCAM-1 exhibited a correspondingly higher presence of LRG1 signals. This evidence strongly suggests a correlation between LRG1 and chronic inflammation associated with atherosclerosis.

### Depleting LRG1 leads to a significant reduction in disease progression in the ApoE deficiency-induced atherosclerotic mouse model

To gain further insights into the pathological mechanism of LRG1, particularly its role in inflammation-related processes during atherosclerosis progression, we utilized a mouse model with ApoE deficiency-induced atherosclerosis and further generated mice with double knockout of *Lrg1* and *Apoe* (Figure S2A).

To characterize this atherosclerotic mouse model, we initially analyzed several baseline characteristics after subjecting the mice to a high-fat diet for 32 weeks. Knocking out *Lrg1* in *Apoe^-/-^* mice led to a reduction specifically in proinflammatory markers, with no observable impact on other measured parameters in plasma. Significantly, the concentrations of total cholesterol, high-density lipoprotein cholesterol (HDL-C), and glucose displayed no statistically significant differences in *Apoe^-/-^/Lrg1^-/-^*mice. Simultaneously, there was a non-significant reduction in body weight, low-density lipoprotein cholesterol (LDL-C), and triglyceride levels in these mice, potentially stemming from a feed-forward effect associated with the attenuated pro-inflammatory state observed throughout the disease course, although this alteration did not attain statistical significance (Figure S2B). Furthermore, consistent with these findings, we observed that the high-fat diet induced an increase in plasma LRG1 levels in *Apoe^-/-^* mice but not in WT or *Apoe^-/-^/Lrg1^-/-^* mice (Figure. S2C), and the ratio of increased plasma LRG1 level of *Apoe^-/-^* mice appeared to rise with age when compared to the relative values from the same aged WT mice (Figure S2D). Collectively, these data strongly suggest that LRG1 contributes to disease progression in the high-fat diet-induced ApoE deficiency-induced atherosclerotic mouse model. We conducted a comprehensive analysis of the size and number of atherosclerotic plaques in the aorta artery at different stages of disease progression, including the onset stage (16 weeks), early stage (25 weeks), progressing stage (29 weeks), and the advanced atherosclerosis stage (32 weeks) in both *Apoe^-/-^ and Apoe^-/-^/Lrg1^-/-^* mice. We employed oil red perfusion to visualize the plaques in the whole aortic artery and observed that knocking out LRG1 from the mice resulted in a significant reduction in the size and number of lesion areas in advanced stage (32 weeks) of atherosclerosis progression (Figure 2A). To further validate this observation, we also examined cross-sections of arteries and the results were consistent with aforementioned findings, showing a significant reduction in the oil-red-positive area when LRG1 was knocked out in *Apoe^-/-^* mice (Figure 2B).

**Figure 2.**
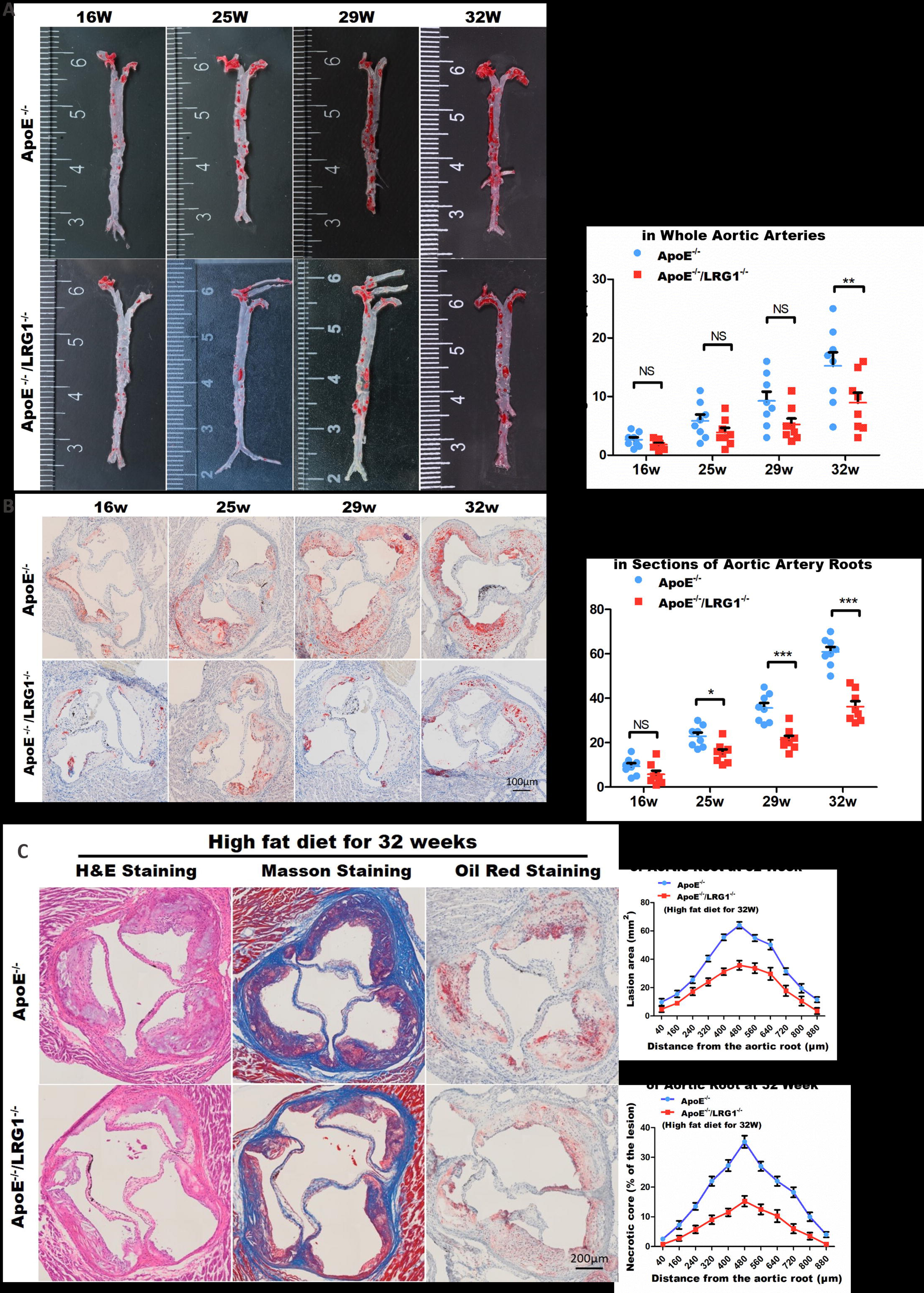
Depleting LRG1 leads to a significant reduction in disease progression in the ApoE deficiency-induced atherosclerotic mouse model. **(A)** Oil Red staining revealed atherosclerotic plaques throughout the entire aortic artery in *Apoe^-/-^* or *Apoe^-/-^ /Lrg1^-/-^* mice following a high-fat diet for 16, 25, 29, or 32 weeks, denoted as 16W, 25W, 29W, and 32W, respectively (depicted on the left). The accompanying chart on the right illustrates the proportions of Oil Red-positive regions in the whole aortic artery across these various groups and time intervals. (**B**) Aortic artery root sections from *Apoe^-/-^* and *Apoe^-/-^/Lrg1^-/-^*mice, fed the high-fat diet for 16W, 25W, 29W, and 32W, were co-stained with H&E and Oil Red. On the left, the images display the staining results. On the right, a plot illustrates the proportions of Oil Red-positive regions in the aortic artery root for each group and time point. (**C**) H&E, Masson, and Oil Red staining were conducted on lesion sections located 480 μm from the aortic artery root in *Apoe^-/-^*and *Apoe^-/-^/Lrg1^-/-^* mice after 32 weeks of a high-fat diet, as shown on the left. On the right, the graphs depict the dynamic changes in the Oil Red-positive lesion area (top) and the percentages of necrotic regions within individual artery sections (bottom). [Mean ± SEM. n.s., not significant, *p < 0.05, **p<0.01, ***p<0.001; One-way ANOVA followed by Tukey’s multiple comparison test]

Particularly, we compared the plaque lesions located 400 μm from the aortic root using oil-red, H&E, and Masson staining at advanced stage (32 weeks) of atherosclerosis progression (Figure 2C). The comparison of plaque lesions between *Apoe^-/-^* and *Apoe^-/-^Lrg1^-/-^* group mice at 32 weeks revealed a consistent reduction in the atherosclerotic lesion area and necrotic core when *Lrg1* knocked out (Figure 2C). Considering all of these findings, it becomes evident that depleting LRG1 leads to a significant reduction in atherosclerosis progression in the ApoE deficiency-induced atherosclerotic mouse model, suggesting that LRG1 plays an important role in promoting atherosclerosis in this mouse model.

### LRG1 contributes to the activation of proinflammatory macrophages in the plasma of the ApoE deficiency-induced atherosclerotic mice

Macrophages are known to play a role in orchestrating chronic inflammation during the progression of atherosclerosis.^19^ We aimed to determine whether the proinflammatory function associated with LRG1 during atherosclerosis progression relies on its impact on macrophages. To investigate this, we conducted a blood routine examination on peripheral blood samples from *Apoe^-/-^* and *Lrg1^-/-^/Apoe^-/-^*mice. Specifically, we observed a significant reduction in the counts of monocytes in the double knockout mice at 8 weeks (P<0.05), while the values for neutrophils and lymphocytes displayed no significant differences between the *Apoe^-/-^* and *Lrg1^-/-^ /Apoe^-/-^* mice (Figure 3A-D).

**Figure 3.**
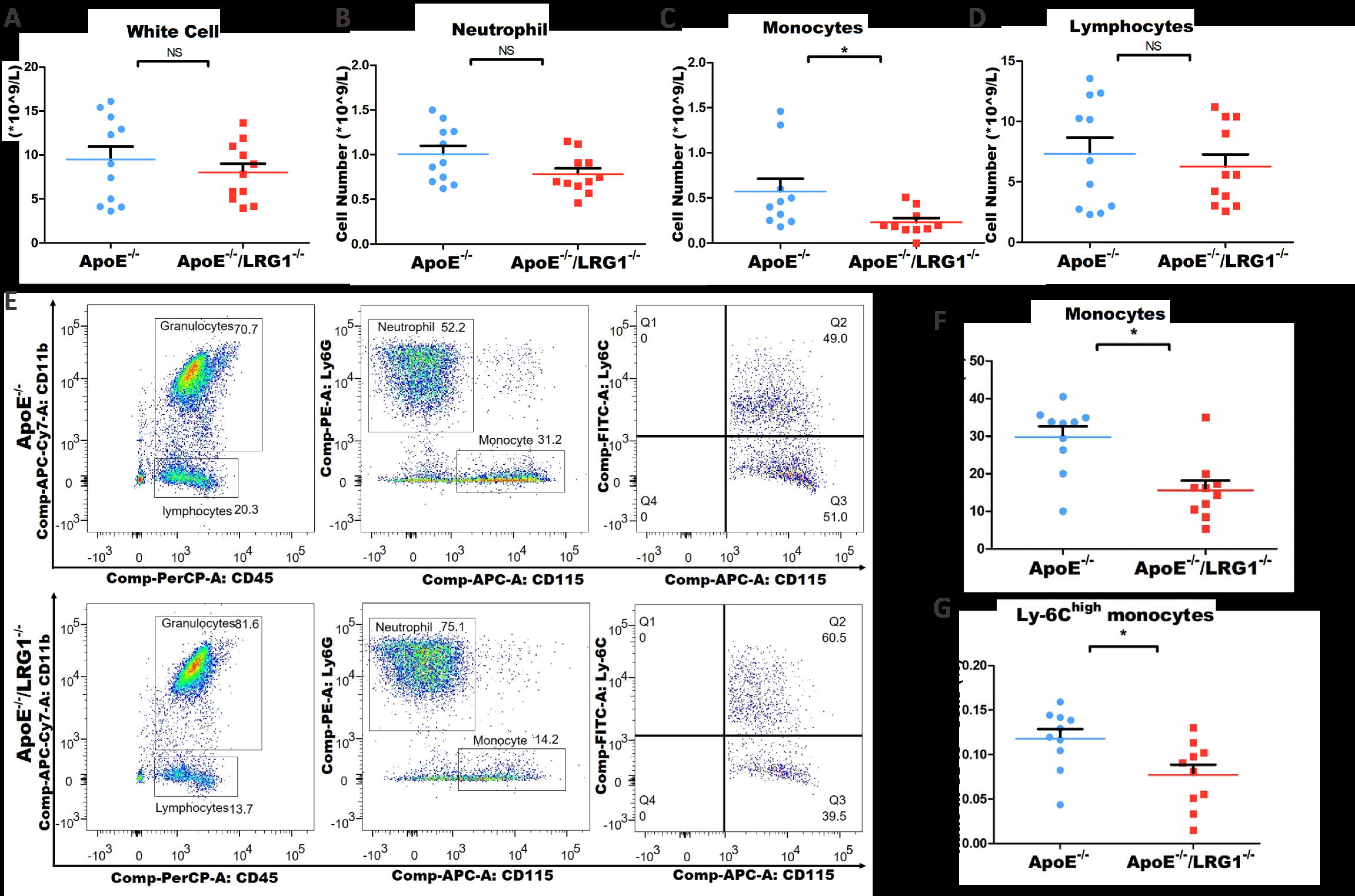
LRG1 contributes to the activation of proinflammatory macrophages in the plasma of the ApoE deficiency-induced atherosclerotic mouse model. **(A-D**) Blood cell composition analysis of *Apoe^-/-^* or *Apoe^-/-^/Lrg1^-/-^* mice following a high-fat diet for 32 weeks: total white blood cell (A), neutrophil (B), monocyte (C), and lymphocyte (D). (**E**) Cell flow cytometry lineage separation in *Apoe^-/-^* (top) and *Apoe^-/-^/Lrg1^-/-^*mice (bottom) fed a 32-week high-fat diet: Left: Ratios of granulocytes (CD11b^+^CD45^+^ cells) or lymphocytes (CD11b^-^CD45^+^ cells) among CD45^+^ white blood cells. Middle: Ratios of neutrophils (Ly6G+CD115^-^CD45^+^ cells) or monocytes (Ly6G^-^CD115^+^CD45^+^ cells) among CD45^+^ white blood cells. Right: Ratios of Ly6C+CD115+CD45+ monocytes or Ly6C^-^CD115^+^CD45^+^ monocytes among total CD115^+^CD45^+^ monocytes. (**F**) The plot contrasts the ratios of monocytes (Ly6G^-^ CD115^+^CD45^+^ cells) within CD45^+^ white blood cells between *Apoe^-/-^* and *Apoe^-/-^ /Lrg1^-/-^* mice following a 32-week high-fat diet. (G) The plot contrasts the ratios of Ly6C^+^ monocytes (Ly6G^-^Ly6C^+^CD115^+^CD45^+^ cells) within CD45^+^ white blood cells between *Apoe-/-* and *Apoe^-/-^/Lrg1^-/-^* mice following a 32-week high-fat diet. [Mean ± SEM. n.s., not significant, *p<0.05, **p<0.01, ***p<0.001; One-way ANOVA followed by Tukey’s multiple comparison test]

Given that a majority of atherosclerotic proinflammatory macrophages originate from blood proinflammatory monocytes (distinguished by their high expression of the Ly-6C cell marker),^2^ we proceeded to compare the ratio of pro-inflammatory to anti-inflammatory monocytes across all peripheral blood cells using flow cytometry. Our findings were consistent with the blood routine examination results, showing a specific and statistically significant reduction (P<0.05) in Ly-6C^High^ (pro-inflammatory monocytes) when *Lrg1* was knocked out from *Apoe^-/-^* mice in the preclinical stage of the atherosclerotic mouse model (Figure 3E-G). We further labeled multiple lineages of plasma cells (Figure S3A) and found that the proportion of neutrophils and lymphocytes, B cells, and T cells displayed no significant differences between the *Apoe^-/-^* and *Lrg1^-/-^/Apoe^-/-^* mice (Figure S3B-D). These results suggest that knocking out *Lrg1* specifically leads to reduced proportions of atherosclerotic proinflammatory monocytes/macrophages in the plasma, consequently delaying the progression of atherosclerosis in a mouse model induced by ApoE deficiency.

### Knocking out *Lrg1* attenuates the accumulation of macrophage-associated proinflammatory cytokines in the plasma and lesions in the ApoE deficiency-induced atherosclerotic mice

To verify the impact of LRG1 deficiency on peripheral proinflammatory monocyte/macrophage cell numbers and their activation, we assessed 23 cytokines using bio-pex (a commercial assay for multi-cytokine analysis per sample). Knocking out *Lrg1* from *Apoe^-/-^* mice notably decreased cytokines released by proinflammatory monocyte/macrophages including IL1α, IL1β, TNF-α, and G-CSF (Figure S4 A-D), as well as those stimulated by proinflammatory macrophages (IFN-γ and IL-6) (Figure S4 E&F) at the disease onset stage of the atherosclerosis mouse model.

We proceeded to assess proinflammatory macrophage activation within aortic plaque sections obtained from both *Apoe^-/-^* and *Lrg1^-/-^/Apoe^-/-^*mice in the advanced stage of atherosclerosis, employing immunohistochemistry. Notably, *Lrg1* knockout in ApoE deficient mice led to a significant reduction in signals for CD68 (a well-established macrophage marker) (Figure 4A), TNF-α (a proinflammatory cytokine released by macrophages) (Figure 4B) and VCAM-1 (a marker of vascular endothelial stress) signals (Figure 4C), highlighting the role of LRG1 in modulating tissue-specific proinflammatory macrophage activation during the disease progression of the atherosclerosis mouse model. The observed attenuation in the activation of tissue-resident proinflammatory macrophages, mediated by LRG1, was accompanied by a consequential decrease in vascular endothelial stress signal VCAM-1, as indicated by its diminished staining. These collective findings suggested that LRG1-regulated macrophage activation could potentially facilitate disease progression in the atherosclerosis mouse model.

**Figure 4.**
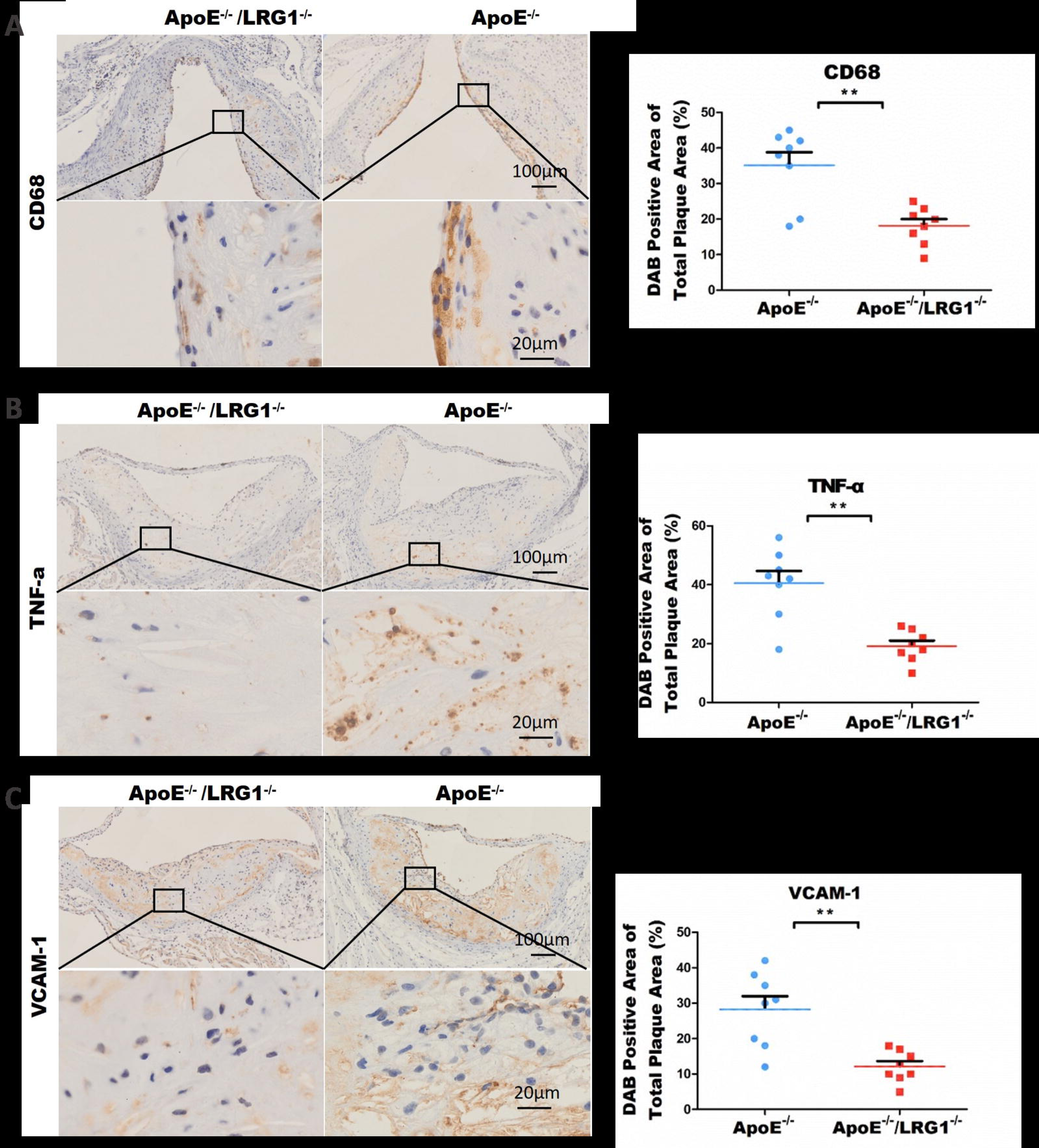
Knocking out LRG1 attenuates the accumulation of macrophage-associated proinflammatory cytokines in plasma and lesions in the ApoE deficiency-induced atherosclerotic mouse model. **(A-C)** The immunohistochemical staining of CD68 **(A)**, TNF-α **(B)**, and VCAM-1 **(C)** on lesion sections located 480 μm from the aortic artery root in *Apoe^-/-^* and *Apoe^-/-^/Lrg1^-/-^*mice after 32 weeks of a high-fat diet. The images on the left display the staining, while the plots on the right illustrate the ratio of DAB-positive area for these targets within the entire detected lesion field. [Mean ± SEM. n.s., not significant, *p<0.05, **p<0.01, ***p<0.001; One-way ANOVA followed by Tukey’s multiple comparison test]

### Treatment with a neutralizing antibody against LRG1 effectively inhibits LRG1-induced macrophage activation and provides significant therapeutic benefits to the ApoE deficiency-induced atherosclerotic mice

To further validate the role of LRG1 in promoting pro-inflammatory macrophage activation, we generated an anti-LRG1neutralizing antibody using recombinant mouse LRG1 (mLRG1) protein to select against a human antibody phage-display library.^20^ After two rounds of library selection, we identified several potent neutralizing antibody candidates (including E4C12, E4D8, E3G1, E3C8, and E4A1) that exhibited high affinity for mLRG1 (Figure 5A and Figure S5B). To confirm their neutralizing activities, we employed an *in vitro* macrophage polarization assay using bone marrow-derived macrophages (BMDMs). Among all candidates, E4C12 displayed the strongest neutralization activity by effectively inhibiting LRG1-induced BMDM polarization, as evidenced by BMDM activation ratio analysis (Figure 5B and Figure S5C).

**Figure 5.**
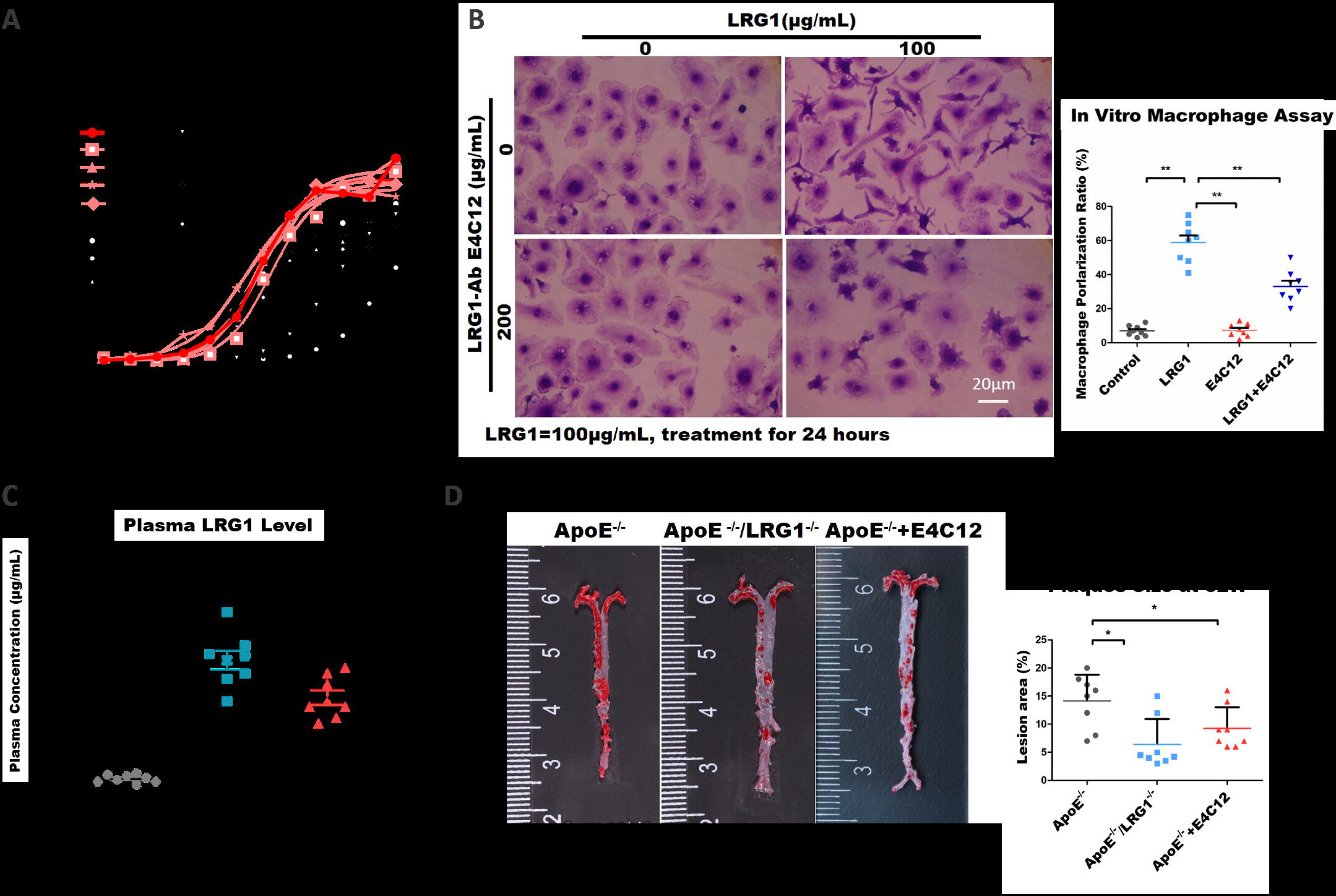
Treatment with a neutralizing antibody against LRG1 effectively inhibits LRG1-induced macrophage activation and provides significant benefits to animals in the ApoE deficiency-induced atherosclerotic mouse model. **(A)** The binding activities of anti-mLRG1 antibodies to mLRG1 analyzed using ELISA. The red lines highlight five anti-mLRG1 antibody candidates that exhibit a high binding activity for mLRG1. **(B)** The images on the left depict in vitro cultured BMDM polarization after 24 hours of treatment with m-LRG1, with and without the m-LRG1 neutralizing antibody E4C12. The plot on the right displays the percentage changes in mLRG1 induced macrophage polarization, both with and without the presence of the mLRG1 neutralizing antibody E4C12. **(C)** Plasma LRG1 levels in *Apoe^-/-^*, *Apoe^-/-^ /LRG1^-/-^*, and *Apoe^-/-^* + E4C12 mice after 32 weeks of a high-fat diet. **(D)** Oil Red staining revealed atherosclerotic plaques throughout the entire aortic artery in *Apoe^-/-^*, *Apoe^-/-^/Lrg1^-/-^*, and *Apoe^-/-^* + E4C12 mice after 32 weeks of a high-fat diet. The images on the left display the staining, while the percentages of the Oil Red staining-positive lesion areas for the three groups are shown on the right. [Mean ± SEM. n.s., not significant, *p < 0.05, **p<0.01, ***p<0.001; One-way ANOVA followed by Tukey’s multiple comparison test]

To assess the potential of the LRG1-neutralizing antibody E4C12 in mitigating atherosclerosis progression, we intraperitoneally administered E4C12 (25mg/kg, 500μg per mouse) (n=12), or control IgG (n=12), to *Apoe^-/-^* mice subjected to a high-fat diet. The treated animals were sacrificed at 32 weeks after high-fat diet feeding (Figure. S5A). This treatment regimen was initiated from the disease onset stage (16 weeks) and continued for 8 weeks, encompassing disease progress period (24 weeks). Remarkably, E4C12 administration significantly lowered the blood LRG1 levels in *Apoe^-/-^* mice during advanced disease stages (Figure 5C). Notably, treatment with the LRG1-neutralizing antibody E4C12 led to a substantial reduction in both the size and quantity of atherosclerotic aortic plaques in the advanced disease stage, with values approaching those observed upon *Lrg1* knockout in *Apoe^-/-^* mice (Figure 5D).

These outcomes underscore the efficacy of diminishing circulating LRG1 levels as a potent strategy to impede atherosclerosis progression in this murine disease model.

### LRG1 stimulates macrophage polarization towards the proinflammatory M1 phenotype

Atherosclerotic macrophage polarization comprises three classical phenotypes: M0 (fully matured/steady state), M1 (proinflammatory, characterized by TNF-α, IL1α, IL1β release under LPS and IFNγ stimulation), and M2 (anti-inflammatory, marked by phagocytosis under IL-4 stimulation).^21^ To further study the pathological mechanisms underlying LRG1-driven macrophage activation and polarization, we conducted an comparison of macrophage morphology within the BMDM macrophage subjected to the polarization assay. Induction of primary macrophage polarization toward M1 or M2-like stages was achieved using pro-inflammatory stimuli (LPS+IFNγ) or the anti-inflammatory stimulus (IL-4), respectively.^22, 23^

Our findings demonstrated that LRG1 stimulation resulted in macrophage polarization more akin to M1 macrophages, rather than M0 or M2 macrophages, with a noteworthy increase in the release of M1 macrophage-associated cytokines, including IL-6, TNF-α, MCP-1, IL-1α, IL-1β, and G-CSF in the macrophage-conditioned medium, compared to the non-treated group (Figure S6A). Importantly, this effect was significantly abrogated upon concurrent treatment with the LRG1-neutralizing antibody E4C12 (Figure 6A).

**Figure 6.**
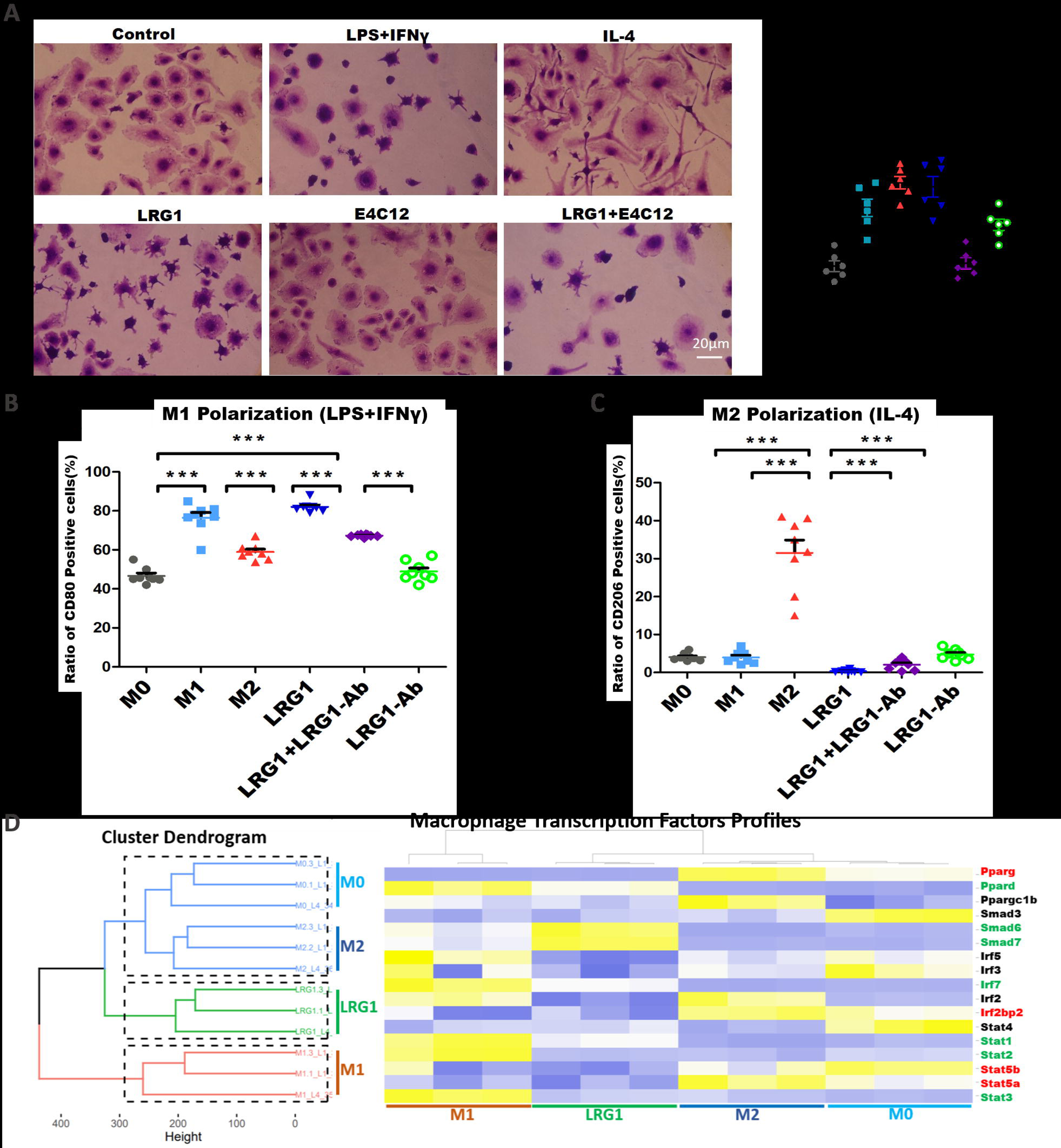
LRG1 stimulates macrophage polarization towards the proinflammatory M1 phenotype. **(A)** The images on the left depict in vitro cultured BMDM polarization after 24 hours of treatment with: DMSO, pro-inflammatory M1-stage stimuli (LPS+IFNγ), the anti-inflammatory M2-stage stimulus (IL-4), mLRG1 with or without the mLRG1 neutralizing antibody E4C12. The plot on the right displays the percentage changes in macrophage polarization after 24 hours treatment above. **(B-C)** The plots illustrate the flow cytometry results, showing the ratios of M1-stage CD80^+^ macrophages **(B)** and M2-stage CD206^+^ macrophages **(C)** among in vitro cultured BMDM after 24 hours of treatment with DMSO, LPS+IFNγ, IL-4, and m-LRG1, both with and without the presence of the m-LRG1 neutralizing antibody E4C12. **(D)** Heatmap depicts the RNA sequencing results, showing the transcription factors profiles of in vitro cultured BMDM after 24 hours of treatment with DMSO, LPS+IFNγ, IL-4, and m-LRG1, both with and without the presence of the m-LRG1 neutralizing antibody E4C12. [Mean ± SEM. n.s., not significant, *p < 0.05, **p<0.01, ***p<0.001; One-way ANOVA followed by Tukey’s multiple comparison test]

To ascertain that LRG1-induced macrophage polarization aligned with M1-like proinflammatory polarization, we evaluated cell surface markers corresponding to the M0 stage (F4/80^+^), M1 stage (F4/80^+^CD80^+^), and M2 stage (F4/80^+^CD206^+^), and stained the macrophages after stimulation with 2 panels of multiple-surface marker staining (Figure S6 B&C). Our analysis revealed that LRG1 treatment induced macrophage polarization characterized by a higher F4/80^+^CD80^+^ ratio akin to M1-polarized macrophages and a reduced F4/80^+^CD206^+^ ratio resembling M2-polarized cells (Figure 6 B&C). The M1-like polarization of macrophage under LRG1 treatment could be neutralized by concurrent treatment with the LRG1-neutralizing antibody E4C12 (Figure S6 B&C).

To further validate the M1-like polarization induced by LRG1, we investigated the activation of macrophage-specific transcription factors associated with the M1 stage. We analyzed a heatmap derived from transcription factor data obtained through RNA sequencing of M0, M1, M2, LRG1-treated, and LRG1+E4C12-treated macrophages. This analysis consistently demonstrated that the LRG1 treatment led to an increase in the expression of M1-specific transcription factors (e.g., Pparg, Irf2bp2, Stat5a, and Stat5b). Conversely, the expression of M2-specific transcription factors (such as Ppard, Smad6, Smad7, Irf7, Stat1, Stat2, and Stat3) remained unaffected (Figure 6D). The complete list of macrophage transcription factors was shown in Figure S6D. Taken together, these findings underscore the ability of LRG1 to induce macrophage M1 stage polarization, a process effectively ameliorated by its neutralizing antibody.

### LRG1-induced macrophage proinflammatory polarization is regulated by the ERK and JNK signaling pathways

Previous studies have discovered that macrophage proinflammatory polarization (M1 stage) triggered by LPS and IFN-r stimulation is primarily regulated by the MAPK and NF-κB signaling pathways.^21^ To investigate whether the same signaling pathway was also utilized by LRG1, we assessed levels of phosphorylated MAPK and NF-κB signaling in mouse BMDMs subjected to LRG1 treatment. Intriguingly, within 1 hour of LRG1 treatment, there was a notable increase in the phosphorylation levels of extracellular signal-regulated kinase 1/2 (ERK1/2), p38, c-Jun as visualized by western blotting analysis (Figure 7A). These results suggested that the activation of ERK1/2, JNK, p38 and p65 potentially mediated the LRG1-induced macrophage M1-like polarization. To substantiate this finding, we employed inhibitors targeting ERK1/2 (AZD6244), JNK (SP600125), p38 (BIRB796) and NF-κB (TCPA-1) and co-treated those inhibitors with LRG1. After confirming the inhibitory efficacy of these inhibitors (Figure S7A-D), we noticed that inhibition of ERK1/2 or JNK partially delayed LRG1-induced macrophage polarization within a 24-hour time frame. Remarkably, concurrent inhibition of ERK1/2 and JNK almost completely abrogated LRG1-induced macrophage polarization within 24 hours (Figure 7B), while inhibiting p38 or NF-κB signaling only mildly influenced LRG1 induced macrophage M1-like polarization (Figure S7E). Taken together, these findings strongly suggest that LRG1-induced macrophage M1-stage polarization is primarily governed by the ERK1/2 and JNK pathways.

**Figure 7.**
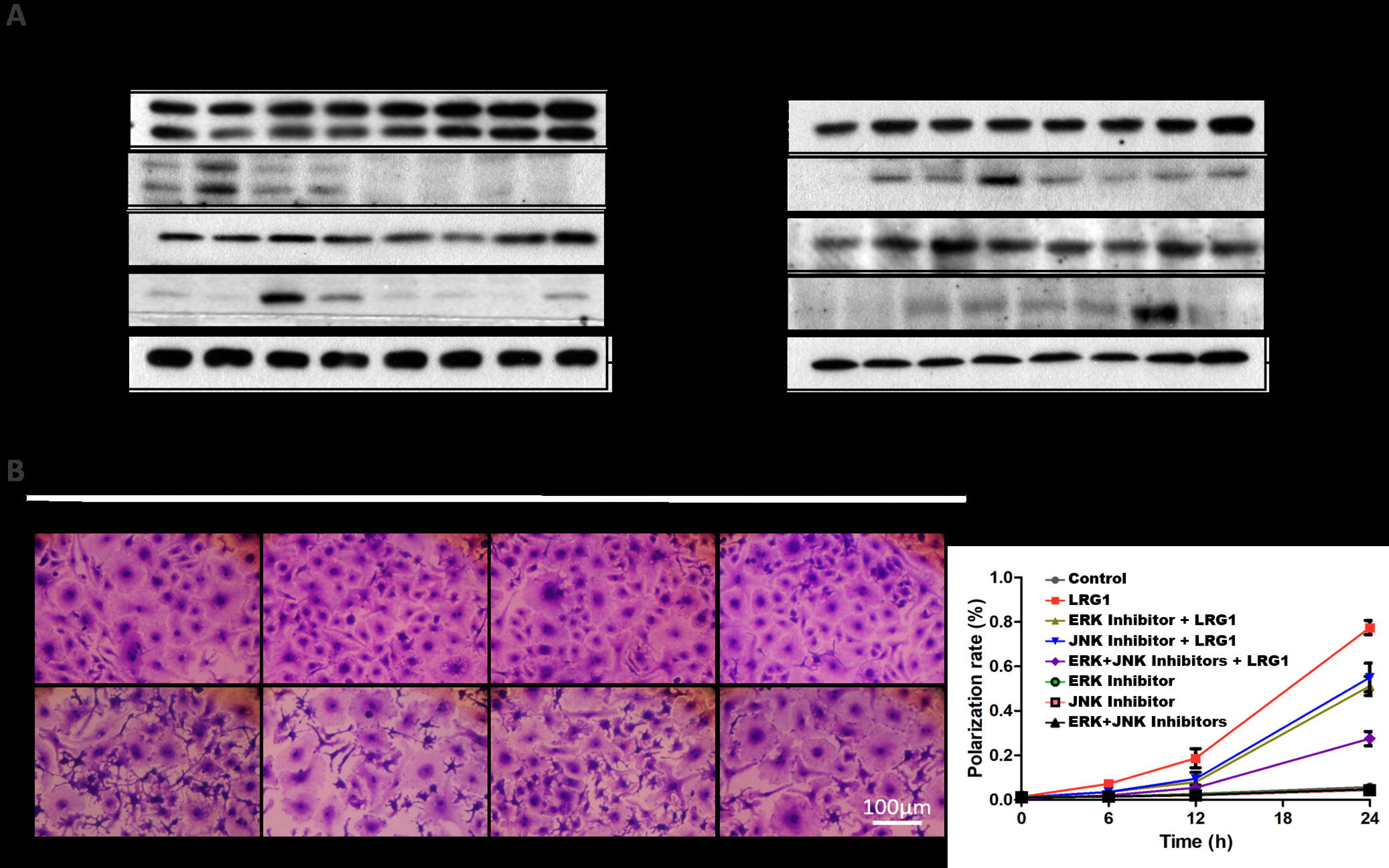
LRG1-induced macrophage proinflammatory polarization is regulated by the ERK and JNK signaling pathways in vitro. **(A)** Immunoblotting detects markers of the activation of MAPK (ERK, JNK, and p38) and NF-κB signaling pathways in samples from in vitro cultured BMDM after 24 hours of LRG1 treatment. The antibodies and phosphorylation-specific antibodies recognize ERK1/2, c-Jun, p38, and p65, reflecting the activation of ERK, JNK, p38, and the NF-κB pathway, respectively. GAPDH is shown as the sample loading control. **(B)** The images on the left depict the polarization of in vitro cultured BMDM after 24 hours of treatment with DMSO and inhibitors targeting ERK1/2 (AZD6244), JNK (SP600125), and a combination of both, both with and without LRG1. The curves on the right represent the percentage changes in macrophage polarization during the 0-24 hours of treatment as described above. [Mean ± SEM. n.s., not significant, *p< 0.05, **p<0.01, ***p<0.001; One-way ANOVA followed by Tukey’s multiple comparison test]

## Discussion

The current study presented compelling evidence substantiates the role of LRG1 in accelerating atherosclerosis progression by inducing pro-inflammatory macrophage polarization. LRG1 does so through the Mitogen-Activated Protein Kinase (MAPK) and JNK signaling pathway within macrophages, resulting in alterations to Toll-like Receptor 4 (TLR4) signaling and an elevation in Nuclear Factor-Kappa B (NF-κB)-mediated proinflammatory gene expression (Figure 7). Of notable significance is the observation that neutralization of plasma LRG1 effectively hinders M1-like macrophage polarization to concurrently reduce atherosclerotic lesion area and diminish aortic inflammation. Therefore, inhibition of LRG1 emerges as a promising novel strategy for combating atherosclerotic cardiovascular disease.

LRG1 is a highly conserved member of the leucine-rich repeat protein family, a group often implicated in protein–protein interactions, cell adhesion, and signaling mechanisms. This protein is primarily localized in the vasculature and exhibits basal expression under normal physiological conditions.^8^ Previous studies have underscored the potential of LRG1 as a significant predictor for CAD patients with familial hypercholesterolemia as well in arterial stiffness, endothelial function, and peripheral artery disease.^18,24–25^ However, due to the inherent limitations of human studies, the prior work have not ascertained the causal impact or predictive potential of LRG1 levels. The current study not only demonstrated that LRG1 is significantly elevated in CAD patients and exhibits correlations with inflammatory factors like high-sensitivity C-reactive protein (hsCRP) and Growth Differentiation Factor 15 (GDF-15), but also indicated its causal role in atherosclerosis using animal models and a neutralizing antibody.

Our LRG1-deficient mice exhibited a potential reduction in atherosclerosis, accompanied by decreased serum lipids when compared to *Apoe* knockout mice. Despite the unclear molecular link between the induction of lipid inflammatory factors in LRG1 knockout mice, necessitating further studies, a significant and persistent disparity in lesion size between the two genotypes was evident. This contrast was particularly pronounced in the aortic arch and root during the advanced stages of the disease. Importantly, inflammatory factors such as Tumor Necrosis Factor-alpha (TNF-α), Interleukin-1 beta (IL-1β), and Interferon-gamma (IFN-γ) showed substantial reductions, specifically in the early stage of atherosclerosis, suggesting a potential plateau phase in lesion growth. This suggests a differential plaque progression at distinct stages. Lesion composition analyses provided insights into decreased macrophage infiltration, concomitant with a significant downregulation of proinflammatory factors like TNF-α and Vascular Cell Adhesion Molecule-1 (VCAM-1). Furthermore, LRG1 deficiency correlated with reduced levels of Ly-6C high monocytes in the blood, indicative of a mitigation in innate immunity implicated in atherogenesis. Notably, we extended our investigation by assessing the impact of neutralizing LRG1 antibodies on established mouse atherosclerosis, thus closely simulating the clinical application of anti-LRG1 therapies for cardiovascular diseases. Encouragingly, neutralization of LRG1 significantly reduced the area occupied by established lesions in the aortic arch and roots.

Inflammation plays a pivotal protagonist in atherogenesis, particularly during the acute phase.^26,27^ Macrophages, acting as both sources and regulators of inflammation, adapt to dynamic changes throughout different stages of atherosclerotic plaque development.^28–30^ As atherosclerosis progression, proinflammatory stimuli spur the migration of hematopoietic stem and progenitor cells from the bone marrow to the spleen. In the spleen, these cells differentiate into monocytes, subsequently released into circulation and recruited to inflamed regions, such as the atherosclerotic vessel wall.^31^ Our study presented here indicated that LRG1 drives macrophages differentiation toward M1 phenotype, thus contributing to inflammation and atherosclerosis.

Our study confirmed that blockade of JNK and ERK prevented the LRG1-induced M1-like macrophage polarization. It is well known that the MAPK pathway regulates the production of inflammatory mediators and controls processes related to atherosclerosis. It is noteworthy that the suppression of JNK phosphorylation by JNK inhibitor SP600125 has been found to promote macrophage shift to an anti-inflammatory phenotype.^32,33^ Similar effects could be found by blocking ERK signaling through the selective inhibitor ERK inhibitor AZD6244.^34^ Importantly, combination of SP600125 and AZD6244 significantly inhibits the macrophage polarization to M1-like phenotype (Figure 7B). In contrast, p38 MAPK inhibitor BIRB 796 and NF-κB inhibitor TCPA-1 did not shift the macrophage polarization. Thus, LRG1 exerts a pro-inflammatory effect by promoting activation of the pathway in macrophages through the enhancement of JNK/ERK signaling.

Atherosclerosis progression and plaque remodeling are determined by the crosstalk between vascular cells and macrophages in advanced plaques. The fact that LRG1 promotes macrophages polarization toward the M1 phenotype underscores the predominant role of LRG1 in inflammation during the atherosclerosis process. It is thus a plausible strategy to further advance the LRG1 neutralizing antibody toward more pre-clinical studies for eventual clinical investigation.

## Supplemental Material

Expanded Methods

Figures S1–S7

Legends for Figures S1–S7

References 1–4

## Reference

1. Libby P. The changing landscape of atherosclerosis. Nature 2021;592:524–533.

2. Tabas I, Lichtman AH. Monocyte-Macrophages and T Cells in Atherosclerosis. Immunity 2017;47:621–634.

3. Weber C, Habenicht AJR, von Hundelshausen P. Novel mechanisms and therapeutic targets in atherosclerosis: inflammation and beyond. Eur Heart J 2023;44:2672–2681.

4. Vergallo R, Crea F. Atherosclerotic Plaque Healing. N Engl J Med 2020;383:846–857.

5. Tabas I, Bornfeldt KE. Macrophage Phenotype and Function in Different Stages of Atherosclerosis. Circ Res 2016;118:653–667.

6. Yurdagul A, Jr. Crosstalk Between Macrophages and Vascular Smooth Muscle Cells in Atherosclerotic Plaque Stability. Arterioscler Thromb Vasc Biol 2022;42:372–380.

7. Childs BG, Baker DJ, Wijshake T, Conover CA, Campisi J, van Deursen JM. Senescent intimal foam cells are deleterious at all stages of atherosclerosis. Science 2016;354:472–477.

8. Barrett TJ. Macrophages in Atherosclerosis Regression. Arterioscler Thromb Vasc Biol 2020;40:20–33.

9. Wang X, Abraham S, McKenzie JAG, Jeffs N, Swire M, Tripathi VB, Luhmann UFO, Lange CAK, Zhai Z, Arthur HM, Bainbridge J, Moss SE, Greenwood J. LRG1 promotes angiogenesis by modulating endothelial TGF-β signalling. Nature 2013;499:306–311.

10. Camilli C, Hoeh AE, De Rossi G, Moss SE, Greenwood J. LRG1: an emerging player in disease pathogenesis. J Biomed Sci 2022;29:6.

11. Fujimoto M, Serada S, Suzuki K, Nishikawa A, Ogata A, Nanki T, Hattori K, Kohsaka H, Miyasaka N, Takeuchi T, Naka T. Leucine-rich α2 -glycoprotein as a potential biomarker for joint inflammation during anti-interleukin-6 biologic therapy in rheumatoid arthritis. Arthritis Rheumatol 2015;67:2056–2060.

12. Hisata S, Racanelli AC, Kermani P, Schreiner R, Houghton S, Palikuqi B, Kunar B, Zhou A, McConn K, Capili A, Redmond D, Nolan DJ, Ginsberg M, Ding BS, Martinez FJ, Scandura JM, Cloonan SM, Rafii S, Choi AMK. Reversal of emphysema by restoration of pulmonary endothelial cells. J Exp Med 2021;218:e20200938.

13. Ha YJ, Kang EJ, Lee SW, Lee SK, Park YB, Song JS, Choi ST. Usefulness of serum leucine-rich alpha-2 glycoprotein as a disease activity biomarker in patients with rheumatoid arthritis. J Korean Med Sci 2014 Sep;29(9):1199–204.

14. Yang Y, Luo R, Cheng Y, Liu T, Dai W, Li Y, Ge S, Xu G. Leucine-rich α2-glycoprotein-1 upregulation in plasma and kidney of patients with lupus nephritis. BMC Nephrol 2020;21:122.

15. Watson CJ, Ledwidge MT, Phelan D, Collier P, Byrne JC, Dunn MJ, McDonald KM, Baugh JA. Proteomic analysis of coronary sinus serum reveals leucine-rich α2-glycoprotein as a novel biomarker of ventricular dysfunction and heart failure. Circ Heart Fail 2011;4:188–197.

16. Chen JH, Chang YW, Yao CW, Chiueh TS, Huang SC, Chien KY, Chen A, Chang FY, Wong CH, Chen YJ. Plasma proteome of severe acute respiratory syndrome analyzed by two-dimensional gel electrophoresis and mass spectrometry. Proc Natl Acad Sci U S A 2004;101:17039–17044.

17. Jiang WJ, Xu CT, Du CL, Dong JH, Xu SB, Hu BF, Feng R, Zang DD, Meng XM, Huang C, Li J, Ma TT. Tubular epithelial cell-to-macrophage communication forms a negative feedback loop via extracellular vesicle transfer to promote renal inflammation and apoptosis in diabetic nephropathy. Theranostics 2022;12:324–339.

18. Pek SL, Tavintharan S, Wang X, Lim SC, Woon K, Yeoh LY, Ng X, Liu J, Sum CF. Elevation of a novel angiogenic factor, leucine-rich-α2-glycoprotein (LRG1), is associated with arterial stiffness, endothelial dysfunction, and peripheral arterial disease in patients with type 2 diabetes. J Clin Endocrinol Metab 2015;100:1586–1593.

19. Canfrán-Duque A, Rotllan N, Zhang X, Andrés-Blasco I. Macrophage-Derived 25-Hydroxycholesterol Promotes Vascular Inflammation, Atherogenesis, and Lesion Remodeling. Circulation 2023;147:388–408.

20. Li D, He W, Liu X, Zheng S, Qi Y, Li H, Mao F, Liu J, Sun Y, Pan L, Du K, Ye K, Li W, Sui J. A potent human neutralizing antibody Fc-dependently reduces established HBV infections. Elife 2017 Sep 26;6:e26738.

21. Van den Bossche J, O’Neill LA, Menon D. Macrophage Immunometabolism: Where Are We (Going)? Trends in immunology 2017;38:395–406.

22. Chinetti-Gbaguidi G, Colin S, Staels B. Macrophage subsets in atherosclerosis. Nat Rev Cardiol 2015;12:10–17.

23. Ganta VC, Choi MH, Kutateladze A, Fox TE, Farber CR, Annex BH. A MicroRNA93-Interferon Regulatory Factor-9-Immunoresponsive Gene-1-Itaconic Acid Pathway Modulates M2-Like Macrophage Polarization to Revascularize Ischemic Muscle. Circulation 2017;135:2403–2425.

24. Bos S, Phillips M, Watts GF, Verhoeven AJM, Sijbrands EJG, Ward NC. Novel protein biomarkers associated with coronary artery disease in statin-treated patients with familial hypercholesterolemia. Journal of clinical lipidology 2017;11:682–693.

25. Pek SLT, Cheng AKS, Lin MX, Wong MS, Chan EZL, Moh AMC, Sum CF, Lim SC, Tavintharan S. Association of circulating proinflammatory marker, leucine-rich-α2-glycoprotein (LRG1), following metabolic/bariatric surgery. Diabetes Metab Res Rev 2018;34:e3029.

26. Wang J, Liu J, Guo W, Bai Y, Li H, Chen H, Han L, Lyu L, Xu C, Liu H. Multiple Biomarkers in the Context of Conventional Risk Factors in Patients With Coronary Artery Disease. J Am Coll Cardiol 2017;69:2769–2770.

27. Wang J, Liu H, Sun J, Xue H, Xie L, Yu S, Liang C, Han X, Guan Z, Wei L, Yuan C, Zhao X, Chen H. Varying correlation between 18F-fluorodeoxyglucose positron emission tomography and dynamic contrast-enhanced MRI in carotid atherosclerosis: implications for plaque inflammation. Stroke 2014;45:1842–1845.

28. Moore KJ, Sheedy FJ, Fisher EA. Macrophages in atherosclerosis: a dynamic balance. Nat Rev Immuno. 2013;13:709–721.

29. Karunakaran D, Nguyen MA, Geoffrion M, Vreeken D, Lister Z, Cheng HS, Otte N, Essebier P, Wyatt H, Kandiah JW, Jung R, Alenghat FJ, Mompeon A, Lee R, Pan C, Gordon E, Rasheed A, Lusis AJ, Liu P, Matic LP, Hedin U, Fish JE, Guo L, Kolodgie F, Virmani R, van Gils JM, Rayner KJ. RIPK1 Expression Associates With Inflammation in Early Atherosclerosis in Humans and Can Be Therapeutically Silenced to Reduce NF-κB Activation and Atherogenesis in Mice. Circulation 2021;143:163–177.

30. Koelwyn GJ, Corr EM, Erbay E. Regulation of macrophage immunometabolism in atherosclerosis. Nat Immunol. 2018;19:526–537.

31. Groh L, Keating ST, Joosten LAB, Netea MG, Riksen NP. Monocyte and macrophage immunometabolism in atherosclerosis. Semin Immunopathol. 2018;40:203–214.

32. Nieminen R, Lahti A, Jalonen U, Kankaanranta H, Moilanen E. JNK inhibitor SP600125 reduces COX-2 expression by attenuating mRNA in activated murine J774 macrophages. Int Immunopharmacol. 2006 Jun;6(6):987–96

33. Li L, Sapkota M, Kim SW, Soh Y. Herbacetin inhibits inducible nitric oxide synthase via JNK and nuclear factor-κB in LPS-stimulated RAW264.7 cells. Eur J Pharmacol. 2015;765:115–23.

34. Zhang Q, Le K, Xu M, Zhou J, Xiao Y, Yang W, Jiang Y, Xi Z, Huang T. Combined MEK inhibition and tumor-associated macrophages depletion suppresses tumor growth in a triple-negative breast cancer mouse model. Int Immunopharmacol. 2019;76:105864.

