## supplementary Methods for "LRG1 promotes atherosclerosis by activating macrophages"

**Human study**

The research adhered to a meticulously devised protocol for the collection of human tissue samples, duly sanctioned by the Institutional Review Board of Beijing Chao-Yang Hospital Capital Medical University and Chinese PLA General Hospital (serial number, 2020-ke-3). Prior to enrollment, all participants offered written informed consent. The in vivo human study encompassed a cohort of 440 consecutive patients, comprising 318 individuals afflicted with Coronary Artery Disease (CAD) and 122 subjects devoid of any discernible evidence or historical record of vascular disease, who served as controls. Control subjects were recruited from outpatient clinics of our hospital due to suspected cardiac chest pain or during their annual physical examinations. Coronary angiography was conducted on all patients, and the diagnosis of CAD was made based on visual observation of luminal diameter narrowing ≥50% in a major epicardial coronary artery.

Conducting experiments with human tissue samples was conducted in strict compliance with relevant guidelines and regulations. Each subject underwent comprehensive medical and family history assessments and provided fasting blood samples during their check-up outpatient visits. All subjects underwent normal chest X-ray examinations, carotid artery ultrasound examinations, resting electrocardiograms, and echocardiography assessments. For each subject, serum biochemical measurements were carried out, including lipid profiles, liver and kidney function tests, as well as assessments of LRG1, GDF-15, and hsCRP levels. Plasma samples were procured from two distinct groups: healthy individuals (n=112) and patients with CAD prior to commencing statin treatment (n=318). Blood samples from patients were obtained after an overnight fasting period and subsequently stored at -80°C until analysis. Serum glucose, glycosylated hemoglobin (HbA1c), creatinine, uric acid, and lipid profiles were measured utilizing standard laboratory techniques on a Hitachi 912 Analyzer (Roche Diagnostics, Germany). Plasma LRG1 levels were quantified using a commercially available ELISA kit (Novus Biotechne, NBP2-60577), while hsCRP levels were determined using a high-sensitivity ELISA kit (Biocheck Laboratories, Toledo, OH, USA), and GDF-15 levels were assessed via an ELISA kit (R&D Systems, Minneapolis, MN, USA).

**Human tissue sample management**

Human femoral and carotid artery tissue samples were obtained from patients diagnosed with severe atherosclerosis who were admitted to the Vascular Surgery Department at the Chinese PLA General Hospital. These patients required surgical interventions due to the total occlusion or embolism of their atherosclerotic carotid and femoral arteries, which were characterized by the presence of atherosclerotic plaques. A total of 10 tissue samples were collected from these patients during their surgical vascular grafting procedures.

Additionally, non-atherosclerotic artery samples were acquired from patients. These non-atherosclerotic samples were obtained from surplus segments of femoral or carotid arteries that would have otherwise been discarded. In total, 10 such non-atherosclerotic artery samples were collected.

To ensure the cleanliness of the tissue samples, they were thoroughly washed with buffered saline. Subsequently, a portion of the tissue samples was preserved in paraformaldehyde (PFA) at a temperature of 4 ℃ until they were needed for further analysis. The remaining portions of the tissue samples underwent a series of preparation steps, which included homogenization and sonication. This was carried out in a lysis buffer composed of 20 mmol/L TRIS (pH 7.4), 150 mmol/L NaCl, 1 mmol/L EDTA, 1 mmol/L EGTA, 1% Triton, 0.1% sodium dodecyl sulfate (SDS), and a protease inhibitor cocktail. The resulting lysates were stored at -80 °C until they were ready to be used for subsequent experiments and analyses.

**Animal studies**

C57BL6/J (WT) mice (stock number C001089), *Apoe^-/-^* mice (stock number S-KO-01101), and *Lrg1^-/-^* mice (stock number S-KO-14978) were procured from Cyagen Biosciences Inc. To generate *Lrg1^-/-^/Apoe^-/-^* double knockout mice, *Lrg1^-/-^* mice were crossbred with *Apoe^-/-^* mice, resulting in F1 progeny that carried both LRG1 and ApoE gene mutations (*Lrg1^+/-^/Apoe^+/-^*). The confirmation of the double knockout condition was achieved through genotyping, utilizing specific primers designed for LRG1 and *Apoe^-/-^* genotyping (primer pairs P1/P2 and P3/P4, respectively), as detailed in Supplementary Figure 2A. For subsequent experiments, male *Apoe^-/-^* mice and male *Apoe^-/-^/Lrg1^-/-^* mice, aged between 6 to 8 weeks, were subjected to a high-fat diet composed of 21% fat, 0.2% cholesterol, 23% protein, and 40.5% carbohydrate. This diet was sourced from Beijing Keao Xieli Feed Co., Ltd. in Beijing, China. To maintain consistency and reduce variability in the experimental groups, mice of the same sex and age were employed. All mice were housed individually in ventilated cages within a pathogen-free facility.

Male mice were specifically chosen for the atherosclerosis experiments to minimize variations and conserve the number of animals used. The assignment of mice to experimental groups was done randomly, and the results were analyzed in a blinded manner to ensure impartiality and objectivity. Genotyping procedures were outsourced to Cyagen, detailed described in Supplementary Figure 2A, where polymerase chain reaction (PCR) was employed for the genotyping process.

**Effect of LRG1 neutralization on atherogenesis**

In this study, *Apoe^-/-^* mice were subjected to a high-fat diet for a duration of 8 weeks, reaching a total study duration of 16 weeks. During this period, these mice received subcutaneous injections of mouse anti-mouse monoclonal antibodies of the IgG2a class. These antibodies were specifically designed to selectively neutralize LRG1, a target protein of interest. An isotype-matched control IgG2a antibody, provided by Novartis, was administered to a control group of mice. The anti-LRG1 antibody exerts its function by binding to LRG1, thereby blocking its interaction with its respective receptors in the plasma.

The neutralizing antibody injections were given at a consistent dose of 50mg/kg. Based on pharmacokinetic assessments conducted by the manufacturers, both the anti-LRG1 and control IgG2a antibodies were administered repeatedly through subcutaneous injections once weekly for a total duration of 8 weeks.

All experimental procedures were carried out in strict accordance with the ethical guidelines and protocols approved by the Institutional Animal Care and Use Committee of Beijing Chao-Yang Hospital, Capital Medical University.

To ensure the statistical power and robustness of our atherosclerotic studies, we predefined the sample size criteria as follows: a minimum of 12 mice per group for morphometric analyses and a minimum of 6 or more mice per group for immunostaining analysis. These predetermined sample sizes were determined through prior analysis of the inherent variability observed in our experiments, aiming to detect a 25% difference between groups with a significance level (alpha) set at 0.05 and a statistical power of 80%.

**In vivo experiments and analysis of atherosclerotic plaques in mice**

Throughout the study, mice were regularly weighed on a weekly basis, starting from birth and continuing until the completion of each experiment. At 6-8 weeks of age, the mice were transitioned from a standard chow diet to a high-fat diet, and they remained on this high-fat diet for a total duration of 32 weeks.

Upon the completion of the experimental protocols and prior to any tissue analysis, the mice were humanely euthanized through the administration of carbon dioxide inhalation. Subsequently, the mice were subjected to perfusion with PBS via the left ventricle of the heart. The heart and the proximal ascending aorta up to the bifurcation of the iliac artery were then carefully excised for subsequent analysis.

The aorta specimens were promptly embedded in Optimal Cutting Temperature (O.C.T) compound and frozen for preservation. To analyze the lesion area within the aorta, as well as for other specified analyses, the aortas were longitudinally split, pinned onto flat black silicone plates, and fixed overnight in a 10% buffered formaldehyde solution in PBS. Following fixation, the aortas were stained with oil red O (Sigma Aldrich) for a duration of 16 hours, rinsed briefly with PBS, and digitally photographed under a consistent magnification using a Carl Zeiss Axio Zoom.V16 microscope.

To assess aortic sinus lesions, cryosections were made at a thickness of 10 μm, and these sections were then stained with oil red O and Masson for overnight incubation. The stained sections were imaged, and digital quantification was performed using Axiovision AC software (Carl Zeiss, Axiovert 200).

Furthermore, hematoxylin and eosin staining (H&E, Sigma Aldrich) was conducted to evaluate plaque morphology. Digital image processing and subsequent morphometric analyses were carried out utilizing the Olympus VS120 software.

Serum samples were collected after the mice had undergone a fasting period of 4-6 hours. These serum samples were then subjected to analysis to determine levels of total cholesterol, HDL-C, LDL-C, triglycerides, and glucose, employing established methods as described in previous references^1,2^.

Moreover, the concentrations of mouse plasma TNF-α, IL-1β, IFN-γ, IL-1α, G-CSF, IL-6 were measured using the Bio-Plex ProTM Mouse Cytokine Standard 23-Plex assay kit (BIO-RAD), and mouse plasma LRG1 levels were determined using ELISA kits from MYBioSource (MBS2020910).

**Immunohistochemistry**

We utilized atherosclerotic and non-significant atherosclerosis femoral artery samples, which were sourced from surplus segments of non-atherosclerotic femoral arteries that would otherwise have been discarded. Upon collection, atherosclerotic arteries were immediately processed and dissected into 0.5 cm segments. Those segments displaying macroscopically evident atherosclerotic lesions were fixed in 4% formaldehyde, embedded in paraffin, and employed for histological staining.

In our histological analysis, consecutive tissue sections underwent staining with various antibodies, including anti-LRG1 (Proteintech, 13224-1-AP), anti-VE-cadherin (Servicebio, GB14013-50), anti-VCAM-1 (Servicebio, GB113498, GB113376), anti-CD68 (Servicebio, GB114109, GB113150), and anti-TNF-α (Servicebio, GB13452). Negative controls were established by omitting primary antibodies and applying isotype-matched control immunoglobulins. Subsequent steps involved treatment with the 3',3'-diaminobenzidine (DAB) chromogen and counterstaining with hematoxylin.

All sections were photomicrographed using an Olympus DP50-CU digital camera mounted on an Olympus BX51 microscope, and the images were digitized for analysis. Importantly, our experiments revealed that immunohistochemistry conducted on femoral atherosclerotic specimens provided superior quality and resolution in elucidating pathophysiological characteristics when compared to coronary atherosclerotic specimens with extensive calcifications.

**Flow Cytometry**

Blood samples were obtained through the retro-orbital puncture and collected into heparinized micro hematocrit capillary tubes, following established procedures as previously describe.^3^ White blood cells were resuspended in a solution comprising 1% fetal bovine serum and 1% bovine serum albumin in PBS. To prevent non-specific binding, the cells were blocked with 2 mg/ml of FcgRII/III before undergoing staining with a cocktail of specific antibodies. Subsequently, the cells were subjected to analysis using a BD flow cytometer.

Monocytes were identified based on the expression of CD45^+^CD11b^+^CD115^+^, with further categorization into Ly6-C^high^ and Ly6-C^low^ subsets. Neutrophils were characterized as CD45^+^Ly6-G^high^, while B cells were distinguished by the presence of CD45^+^CD19^+^ markers. T cells were differentiated into CD45^+^CD4^+^ and CD45^+^CD8^+^ populations. All antibodies employed in the flow cytometry experiments were procured from BioLegend.

Primary macrophages were isolated from femurs and tibias, and bone marrow-derived macrophages (BMDM) and peripheral blood were prepared by macerating the samples through a 70-μm cell strainer (BD Falcon). Prior to analysis, Fc receptors were blocked using the CD16/32 monoclonal antibody (BD, clone 2.4G2). The cells were then stained with a range of fluorochrome-conjugated mouse antibodies, including CD45-PerCP, CD11b-APC-Cy7, CD115-APC, Ly6G-PE, Ly6C-FITC, CD19-BV421, CD3e-PE-Cy7, F4/80-APC, CD206-BV421, and CD86-PE, all of which were obtained from BD Biosciences. Cytometric data were acquired using an LSRFortessa flow cytometer (BD Biosciences) and subsequently analyzed using the FlowJo software (Three Star Inc.).

**Primary cell isolation and cell culture**

BMDMs were isolated from the femurs and tibias of WT mice aged 6-10 weeks. These cells were cultured in DMEM supplemented with 10% FBS, 100 U/ml penicillin, and 100 µg/ml streptomycin. Additionally, 10% conditioned media from L929 cells were added to facilitate macrophage differentiation over a 7-day period.

To activate these BMDMs in vitro, they were polarized into distinct states: classically activated macrophages (M1) were induced by treatment with lipopolysaccharide (LPS, 1μg/ml) and IFNγ (Peprotech, AF-315-05, 50ng/ml), while alternatively activated macrophages (M2) were generated by treatment with IL-4 (10ng/mL). Furthermore, macrophages exposed to LRG1 (100μg/mL) were referred to as M (LRG1).

To assess the influence of LRG1 on BMDMs, we conducted cellular experiments wherein we determined the appropriate concentrations of LRG1 based on both serum levels and the fact that tissue concentrations of this adipokine typically surpass those found in the serum. Subsequently, after a 24-hour exposure to LRG1 protein, we assessed the secretion of inflammatory factors, including TNF-α, IL-1β, MCP-1, IL-1α, IL-6, and granulocyte-macrophage colony-stimulating factor (GM-CSF), using the culture medium of BMDMs.

**Western blotting**

Protein concentrations in both human tissue samples and mouse BMDM cell lysates were quantified using the Bradford method. Tissue samples were lysed in RIPA buffer (containing 50 mM Tris-HCl pH 7.4, 150 mM NaCl, 0.1% SDS, 1% NP-40, 0.25% sodium deoxycholate, 1 mM sodium fluoride, and 1 mM Na3VO4), supplemented with an EDTA-free protease inhibitor cocktail (Roche Life Science, 11 873 580 001) and a phosphatase inhibitor cocktail (Roche Life Science, 04 906 837 001). Protein samples were separated by SDS-PAGE, followed by transfer onto polyvinylidene difluoride membranes.

The membranes were blocked using 5% skim milk (BD Difco, 232100) in PBS containing 0.05% Tween-20 (0.05% TBST). Subsequently, they were incubated with primary antibodies at 4°C for 24 hours. After this incubation, the membranes were probed with horseradish peroxidase (HRP)-conjugated secondary antibodies for 1 hour, and protein bands were visualized using an enhanced chemiluminescence (ECL) detection Kit (ORT2655, PerkinElmer, Waltham, MA, USA).

Primary antibodies employed in this study included phospho-ERK (CST, #4370), ERK (CST, #4695S), phospho-P38 (anti-ACTIVE, #v121C), P38 (CST, #9211), phospho-p65 (93H1, CST, #3033S), P65 (D14E12, CST, #8242S), JNK (CST, #9252S), phospho-JNK (CST, #9251S), c-jun (CST, #8242S), phospho-c-jun (CST, #3270S), GAPDH (MBL, 3H12), α-tubulin (C-11; MBL, PM-054), and LRG1 (Santa Cruz, sc517443). Subsequently, appropriate horseradish peroxidase (HRP)-conjugated secondary antibodies were applied (all from EMD Millipore). Immunoreactive bands were visualized using ECL™ western blot reagents (GE Healthcare Life Sciences).

For ELISA analyses, human plasma and mouse plasma, along with the culture medium of cultured mouse macrophages, were assessed using commercially available ELISA kits. LRG1 levels were quantified using the Novus Biological ELISA kit (NBP2-60577), following the manufacturer's instructions. Additionally, ELISA kits for TNF-α, IL-1β, IL-6, MCP-1, G-CSF, and IL-1α were utilized for the measurement of mouse plasma and culture medium components, as per the manufacturer's guidelines.

**RNA sequencing (RNA-seq) analysis**

Total RNA was extracted from macrophages subjected to various treatments: LPS+IFNγ (M1) from WT BMDMs, IL4 (M2) treated macrophages, LRG1-treated macrophages, and macrophages exposed to LRG1-neutralized antibody E4C12. RNA purity and concentration were assessed using the NanoDrop 2000, while RNA integrity and quantity were determined using the Agilent 2100/4200 system. Following library construction, the library's concentration was quantified using the Qubit® platform, and the samples were sequenced on the novase6000 platform (Illumina).

Sequence reads were quantified using StringTie-1.3.044, and the data were normalized using fragments per kilobase of transcript per million mapped reads (FPKM). Subsequent analyses, including the generation of heatmaps, were performed using R packages (ver. 3.4.1). Differential gene expression analysis was conducted using DeSeq238, which employs negative linearized binomial modeling to estimate dispersion and identify differentially expressed genes in M1 (LPS/IFNγ), M2 (IL-4), LRG1-treated, and LRG1-neutralized antibody (E4C12) macrophages.

In brief, genes with a mean expression of at least 50 counts were considered as expressed, and those showing a minimum 1.5-fold change (with a false discovery rate <0.05) were classified as differentially expressed. Pathway enrichment analysis of the differentially expressed mRNAs was performed using Ingenuity Pathway Analysis (IPA) (Qiagen), with significance determined by a 1-sided Fisher exact test (FDR-adjusted p-value <0.05 and absolute z score >2).

**Expression and purification of recombinant proteins**

The mouse LRG1 (mLRG1) protein spanning residues Leu33 to Leu342 was generated as a His6-Avi-tagged fusion protein, denoted as mLRG1-His6-Avi. This fusion protein was produced through transient transfection of FreeStyle 293F cells and subsequently purified using Ni bead affinity chromatography.

For the full-length IgG antibodies, the coding sequences of the variable heavy chain (VH) and variable light chain (VL) were subcloned into separate expression vectors, one for the mouse IgG1 heavy chain (HC) and the other for the light chain (LC). These expression vectors for HC and LC were introduced into 293F cells via co-transfection at a 1:1 ratio. After 3–6 days of transfection, the cell culture supernatants were harvested for the purification of IgG1 antibodies using Protein A bead affinity chromatography.

**Antibody library panning and screening of anti-LRG1 neutralized antibodies**

To generate anti-mLRG1 antibodies, the mLRG1 fusion protein (mLRG1-His6-Avi) was biotinylated using BirA ligase, resulting in mLRG1-His6-Avi-Biotin. Subsequently, mLRG1-His6-Avi-Biotin was employed as the antigen in panning experiments with a human non-immune antibody library^4^, which had a library size of 1.1x10^10. After two rounds of selection, phage-single-chain variable fragments (phage-scFvs) were screened to identify those exhibiting specific binding to mLRG1.

Approximately 480 individual clones were randomly chosen and subjected to screening for their binding affinity to mLRG1 using an enzyme-linked immunosorbent assay (ELISA). From this pool, 25 phage-scFv antibodies with unique sequences were selected and subsequently converted into the full-length mouse IgG1 format. To further refine the selection, 18 mouse IgG1 antibodies that demonstrated superior performance were chosen based on ELISA-based analyses of their binding to mLRG1.

**ELISA-based binding assays**

To assess the binding between antibodies and antigens, biotinylated antigens were immobilized by capturing them with streptavidin-coated 96-well plates (from Sigma-Aldrich or Nunc, MaxiSorpTM plates). Subsequently, serially diluted antibodies were introduced into the wells and their presence was detected using an HRP-labeled goat polyclonal anti-human IgG Fc antibody, sourced from Thermo Fisher. This assay setup allowed for the measurement and quantification of antibody-antigen interactions.

**Supplementary Figure 1.** **LRG1 elevation correlates with heightened macrophage-derived proinflammatory cytokines in femoral plaques of atherosclerotic femoral stenosis patients. (A-C**) The plot shows the LRG1 signal-positive area among intima **(A)**, media **(B)**, and adventitia **(C)** from atherosclerotic or normal artery slides of CAD patients. (**D-G**) The plot shows the chronic atherosclerosis markers signal-positive area among whole regions of femoral stenosis or normal arteries slides from femoral stenosis patients, CD68 **(D)**, VE-Cadherin **(E)**, VCAM-1 **(F)**, LRG1 **(G)**. [Mean ± SEM. n.s., not significant, *p< 0.05, **p<0.01, ***p<0.001; One-way ANOVA followed by Tukey's multiple comparison test]

**Supplementary Figure 2.** **Depleting LRG1 leads to a significant reduction in disease progression in the ApoE deficiency-induced atherosclerotic mouse model**.

**(A**) The graphic illustrates a genotyping strategy for identifying WT, *Lrg1^-/-^*, *Apoe^-/-^*, and *Apoe^-/-^/Lrg1^-/-^* mice, displayed on the left side. The accompanying table on the right presents the primer sequences used for these mice above. (**B**) The table provides an overview of the baseline characteristics of *Apoe^-/-^* and *Apoe^-/-^/Lrg1^-/-^* mice after being fed a high-fat diet for 32 weeks. (**C**) The plot displays the plasma LRG1 levels in WT, *Apoe^-/-^*, and *Apoe^-/-^/Lrg1^-/-^* mice after fed a high-fat diet for 32 weeks. (D) The plot exhibits the elevated plasma LRG1 ratios of *Apoe^-/-^* mice resulting from an 8, 16, and 32-week high-fat diet. [Mean ± SEM. n.s., not significant, *p< 0.05, **p<0.01, ***p<0.001; One-way ANOVA followed by Tukey's multiple comparison test]

**Supplementary Figure 3.** **LRG1 contributes to the activation of proinflammatory macrophages in the plasma of the ApoE deficiency-induced atherosclerotic mouse model. (A**) Cell flow cytometry lineage separation in *Apoe^-/-^* (top) and *Apoe^-/-^/Lrg1^-/-^* mice (bottom) fed a 32-week high-fat diet: Left: Ratios of granulocytes (CD11b^+^CD45^+^ cells) or lymphocytes (CD11b^-^CD45^+^ cells) among CD45^+^ white blood cells. Middle: Ratios of B cells (CD11b^-^CD19^+^CD45^+^ cells) among CD45^+^ white blood cells. Right: Ratios of T cells (CD11b^-^CD3^+^CD45^+^ cells) among CD45^+^ white blood cells. **(B-D)** These plots compare the ratios of lymphocytes (CD11b^-^CD45^+^ cells) in **(B)**, B cells (CD11b^-^CD119^+^CD45^+^ cells) in **(C)**, and T cells (CD11b^-^CD3^+^CD45^+^ cells) in **(D)** among CD45^+^ white blood cells in *Apoe^-/-^* and *Apoe^-/-^/Lrg1^-/-^* mice after a 32-week high-fat diet. [Mean ± SEM. n.s., not significant, *p< 0.05, **p<0.01, ***p<0.001; One-way ANOVA followed by Tukey's multiple comparison test]

**Supplementary Figure 4.** **Knocking out LRG1 attenuates the accumulation of macrophage-associated proinflammatory cytokines in plasma and lesions in the ApoE deficiency-induced atherosclerotic mouse model. (A-F)** These plots depict the results of ELISA analyses for blood proinflammatory monocyte/macrophage-released cytokine levels: IL1α **(A)**, IL1β **(B)**, TNF-α **(C)**, and G-CSF **(D)**, as well as blood proinflammatory macrophages-related cytokine levels: IFN-γ **(E)** and IL-6 **(F)**, in *Apoe^-/-^* and *Apoe^-/-^/Lrg1^-/-^* mice following 32 weeks of a high-fat diet. [Mean ± SEM. n.s., not significant, *p<0.05, **p<0.01, ***p<0.001; One-way ANOVA followed by Tukey's multiple comparison test]

**Supplementary Figure 5. Treatment with a neutralizing antibody against LRG1 effectively inhibits LRG1-induced macrophage activation and provides significant benefits to animals in the ApoE deficiency-induced atherosclerotic mouse model. (A)** The table provides an overview of the baseline characteristics of *Apoe^-/-^*, *Apoe^-/-^/Lrg1^-/-^*, and *Apoe^-/-^*+E4C12 mice after being fed a high-fat diet for 32 weeks. **(B)** The table lists the EC50 values for the binding activity between the anti-mLRG1 antibodies shown inf Figure 5A with the mLRG1 protein. **(C)** The images on the left depict in vitro cultured BMDM polarization after 24 hours of treatment with mLRG1, with and without the m-LRG1 neutralizing antibodies. The plot on the right displays the percentage changes in mLRG1-induced macrophage polarization, both with and without the presence of the mLRG1 neutralizing antibodies. [Mean ± SEM. n.s., not significant, *p< 0.05, **p<0.01, ***p<0.001; One-way ANOVA followed by Tukey's multiple comparison test]

**Supplementary Figure 6. LRG1 stimulates macrophage polarization towards the proinflammatory M1 phenotype. (A)** The curves represent the proinflammatory cytokines release levels over a time range of 0-24 hours during LRG1 treatment of in vitro cultured BMDM. **(B-C)** The graphics illustrate the flow cytometry strategy, showing the ratios of M1-stage CD80^+^F4/80^+^ macrophages **(B)** and M2-stage CD206^+^F4/80^+^ macrophages **(C)** among in vitro cultured F4/80+ BMDM after 24 hours of treatment with DMSO, LPS+IFNγ, IL-4, and mLRG1, both with and without the presence of the mLRG1 neutralizing antibody E4C12. **(D)** The table lists the RNA sequencing results, showing both the M1- and M2-stage macrophage related transcription factors after 24 hours of treatment with DMSO, LPS+IFNγ, IL-4, and mLRG1, both with and without the presence of the mLRG1 neutralizing antibody E4C12. [Mean ± SEM. n.s., not significant, *p< 0.05, **p<0.01, ***p<0.001; One-way ANOVA followed by Tukey's multiple comparison test]

**Supplementary Figure 7. LRG1-induced macrophage proinflammatory polarization is regulated by the ERK and JNK signaling pathways in vitro. (A-D)** Immunoblotting detects markers of the activation of MAPK (ERK, JNK, and p38) and NF-κB signaling pathways in samples from in vitro cultured BMDM from 0-24 hours of LRG1 treatment, both with or without inhibitors of: the ERK pathway **(A)**, the JNK pathway **(B)**, the p38 pathway **(C)**, and the NF-κB pathway **(D)**. The antibodies and phosphorylation-specific antibodies recognize ERK1/2, c-Jun, p38, and p65, reflecting the activation of ERK, JNK, p38, and the NF-κB pathway, respectively. GAPDH is shown as the sample loading control. **(E)** The images on the left depict the polarization of in vitro cultured BMDM after 24 hours of treatment with DMSO and inhibitors targeting p38 (BIRB796), NF-κB (TCPA-1) and a combination of both, both with and without LRG1. The curves on the right represent the percentage changes in macrophage polarization during the 0-24 hours of treatment as described above. [Mean ± SEM. n.s., not significant, *p< 0.05, **p<0.01, ***p<0.001; One-way ANOVA followed by Tukey's multiple comparison test]

**References**

1. Wang J, Liu J, Guo W, Bai Y, Li H, Chen H, Han L, Lyu L, Xu C, Liu H. Multiple Biomarkers in the Context of Conventional Risk Factors in Patients With Coronary Artery Disease. *J Am Coll Cardiol* 2017;69:2769-2770.

2.	Wang J, Han LN, Ai DS, Wang XY, Zhang WJ, Xu XR, Liu HB, Zhang J, Wang P, Li X, Chen ML. Growth differentiation factor 15 predicts cardiovascular events in stable coronary artery disease. *J Geriatr Cardiol* 2023;20:527-537

3. Tian X, Liu X, Ding J, Wang F, Wang K, Liu J, Wei Z, Hao X, Li Y, Wei X, Zhang H, Sui J. An anti-CD98 antibody displaying pH-dependent Fc-mediated tumour-specific activity against multiple cancers in CD98-humanized mice. *Nat Biomed Eng* 2023;7:8-23

4. Li D, He W, Liu X, Zheng S, Qi Y, Li H, Mao F, Liu J, Sun Y, Pan L, Du K, Ye K, Li W, Sui J. A potent human neutralizing antibody Fc-dependently reduces established HBV infections. *Elife*. 2017 26;6:e26738
